# Mechanisms of HIV Latency in Hematopoietic Progenitors: GFI1 as a Key Regulator

**DOI:** 10.64898/2026.02.19.706859

**Authors:** Jackline A. Onyango, Chen Li, Cuie Chen, W Miguel Disbennett, Maria C. Virgilio, Mark M. Painter, Valeri H. Terry, Barkha Ramnani, Joshua D. Welch, Kathleen L. Collins

**Affiliations:** Department of Biological Chemistry, University of Michigan, Ann Arbor, MI, USA; Department of Computational Medicine and Bioinformatics, University of Michigan, Ann Arbor, MI, USA; Department of Internal Medicine, University of Michigan, Ann Arbor, MI, USA; Post-Baccalaureate Research Education Program (PREP), University of Michigan, Ann Arbor, MI, USA; Cellular and Molecular Biology Program, University of Michigan, Ann Arbor, MI, USA; Graduate Program in Immunology, University of Michigan, Ann Arbor, MI, USA; Department of Computer Science and Engineering, University of Michigan, Ann Arbor, USA; Department of Microbiology and Immunology, University of Michigan, Ann Arbor, MI, USA

**Keywords:** HIV latency, HSPCs, GFI1, transcriptional repression, scRNA-seq

## Abstract

Durable control of HIV infection is challenged by persistent latent reservoirs, including hematopoietic stem and progenitor cells (HSPCs), which provide a unique niche for proviral silencing. Mechanisms of HIV latency in HSPCs remain poorly studied. Here, we utilized a dual-reporter HIV (89.6 VT1) and single-cell RNA sequencing to identify host factors governing latency in HSPCs. Transcriptomic profiling revealed elevated expression of several genes in latently infected cells, among them the transcriptional repressor GFI1. Functional studies showed that GFI1 suppresses HIV gene expression by binding a conserved sequence near the primer binding site within the long terminal repeat (LTR). Disruption of GFI1 DNA-binding or corepressor recruitment domains diminished its silencing effect. GFI1 also antagonized Tat and NF-κB-mediated activation, and reversal of GFI1-mediated suppression was most robust with combined HDAC inhibition and NF-κB activation. These findings position GFI1 as an important regulator of HIV latency in HSPCs.

## INTRODUCTION

Despite transformative advances in HIV treatment through combination antiretroviral therapy (cART), curing HIV remains elusive due to the challenge presented by latent viral reservoirs^1–3^. While suppressive therapy rapidly inhibits HIV replication and prevents new cellular infections, resulting in a decline in infected cells and viral loads to clinically undetectable levels in treated individuals, the therapy itself does not eliminate all infected cells^4^. Instead, a small pool of long-lived cells harboring an integrated virus, often seeded early during infection, persists in a dormant state^2,5,6^. These reservoirs can reignite systemic infection if therapy is interrupted, necessitating lifelong treatment^1,2^. The inability to eliminate these latently infected cells is the central barrier to a true HIV cure, underscoring the urgent need to unravel the biological mechanisms that sustain HIV persistence.

Although CD4^+^ T cells are recognized as the primary cellular reservoir for latent HIV, additional cell types, such as hematopoietic stem and progenitor cells (HSPCs), also sustain lifelong infection ^7–9^. HSPCs are long-lived, self-renewing cells crucial for generating all blood and immune cells^10^. Their ability to persist in a quiescent state, differentiate into multiple lineages, and regulate susceptibility to apoptosis provides a unique environment for the maintenance of latent HIV^11^. Unlike CD4^+^ T cells, in which HIV latency is established after a period of activity, latency in HSPCs is established with viral integration, circumventing the cytotoxicity caused by viral protein expression^7,8,12,13^. These properties allow infected HSPCs not only to sustain latent infection across their lifespan but also potentially to propagate HIV throughout the hematopoietic compartment by giving rise to diverse infected progeny^14^.

Recent evidence highlights the significant contribution of HSPCs to persistent plasma viremia among individuals on long-term antiretroviral therapy^9^. Though less abundant in the circulation than T cells, HSPCs can be infected by HIV and subsequently transmit integrated proviral genomes to both CD4^+^ and CD4^−^ descendant cells^7,15^. Notably, these infected HSPCs are capable of producing virions upon latency reversal in vitro^13^. Sequence analyses have shown that plasma virus often matches clonally expanded proviral genomes originating from HSPCs, and in some donors, these HSPC-derived clones are a quantitatively dominant source of residual plasma viremia^9^. Thus, the unique biological properties and persistence of HSPCs underline their complex role in sustaining HIV latency and reinforce the necessity of further research into latent infection mechanisms and potential therapeutic strategies targeting these cells.

The distinguishing feature between active and latent HIV infection is the presence of a transcriptionally silent provirus in the host genome. Transcriptional regulation, particularly through chromatin condensation and epigenetic mark establishment, plays a pivotal role in both the maintenance and stability of the latent viral reservoir^2,16–20^. Although HIV preferentially integrates into open chromatin regions, latency is closely associated with subsequent chromatin remodeling and the accrual of repressive modifications, including histone deacetylase (HDAC)-mediated deacetylation, methylation by histone methyltransferases, and DNA methylation at CpG islands near the viral promoter^18,19,21–23^. These modifications restrict the access of transcription factors and RNA polymerase II, presenting formidable barriers to spontaneous reactivation and therapeutic latency reversal. Beyond these global epigenetic changes, locus-specific DNA elements within and adjacent to the retroviral promoter, such as the primer binding site (PBS), further fine-tune local chromatin architecture through interactions with host repressor proteins and corepressor complexes^24–27^. Overcoming the multi-layered repression at the HIV promoter requires the coordinated activation of key transcriptional activators, including the nuclear translocation of NF-κB and the viral transactivator Tat, both of which are essential for productive transcription and reactivation from latency^8,28–34^.

The pursuit of a cure for HIV has increasingly focused on strategies that promote reactivation of the virus from latently infected cells, aiming to eliminate these persistent reservoirs. Despite the advances, most studies investigating HIV latency mechanisms have centered on CD4^+^ T cells, leaving the regulatory landscape of other cell types, such as HSPCs, largely underexplored. Earlier evidence suggests that latency in HSPCs may be regulated differently than in T cells, but the specific molecular players, transcriptional context, and chromatin environments responsible for these potential differences have not yet been fully defined^12^. Without a detailed understanding of these processes in HSPCs, interventions designed to target latent reservoirs risk overlooking critical cellular diversity and thereby may fall short of achieving lasting remission.

Notably, there are marked differences in the reactivation response between HSPCs and T cells. For example, Painter et al. demonstrated that despite efficiently increasing histone acetylation in HSPCs^12^, latent provirus in these cells is resistant to viral reactivation, unlike the robust reactivation observed in T cell models^35,36^. Likewise, HMBA, which is a potent latency reversal agent in T cells^37,38^, failed to produce comparable reactivation in quiescent HSPCs^12^. This resistance may be linked to the higher basal nuclear levels of P-TEFb found in unstimulated HSPCs compared to T cells^8^. The heterogeneity in latency reversal agent (LRA)-mediated reactivation across cell types underscores the need to identify the unique molecular mechanisms underlying latency and reactivation in different cellular contexts.

To address critical gaps in our understanding of HIV latency in HSPCs, this study employed a sensitive HIV reporter system (89.6 VT1) to transduce HSPCs and systematically identify host factors associated with latent infection. Leveraging single-cell RNA sequencing (scRNA-seq), we observed distinct transcriptional profiles between latently and actively infected HSPCs, notably the upregulation of the transcriptional repressor GFI1 in the latent state. Functional in vitro assays revealed that GFI1 suppresses HIV gene expression by directly targeting a sequence within the 5’ viral long terminal repeat (LTR) adjacent to the primer binding site and recruiting corepressor complexes to enforce transcriptional silencing. Importantly, GFI1 antagonizes HIV-activating transcription factors such as Tat and NF-κB, thereby helping to maintain latency. Our results further demonstrated that reversing GFI1-mediated repression, using a combination of histone deacetylase inhibition and NF-κB activation, effectively restores HIV gene expression. These findings not only elucidate a novel mechanism of HIV silencing but also advance our knowledge of targeted approaches aimed at eliminating diverse reservoirs of latent HIV.

## RESULTS

### The HIV Reporter VT1 enables the identification of latently and actively infected HSPCs

It is well established that two nucleosomes are consistently positioned within the HIV 5’ long terminal repeat (LTR) of integrated provirus, and multiple host-mediated mechanisms, including recruitment of chromatin-modifying enzymes, have been associated with the establishment of HIV latency^39–42^. However, HIV gene expression may be silenced by host factors and pathways that vary depending on cell type and cell state (Fig. 1A). Here, we sought to identify host factors involved in silencing HIV gene expression in hematopoietic stem and progenitor cells (HSPCs). We utilized the HIV molecular clone 89.6, which our lab modified to create VT1 (Fig. 1B)^34^. VT1 expresses mCherry under the control of the native HIV LTR to monitor active infection and enhanced green fluorescent protein (eGFP) under the spleen focus-forming virus (SFFV) promoter to label all infected cells.

**Fig. 1:**
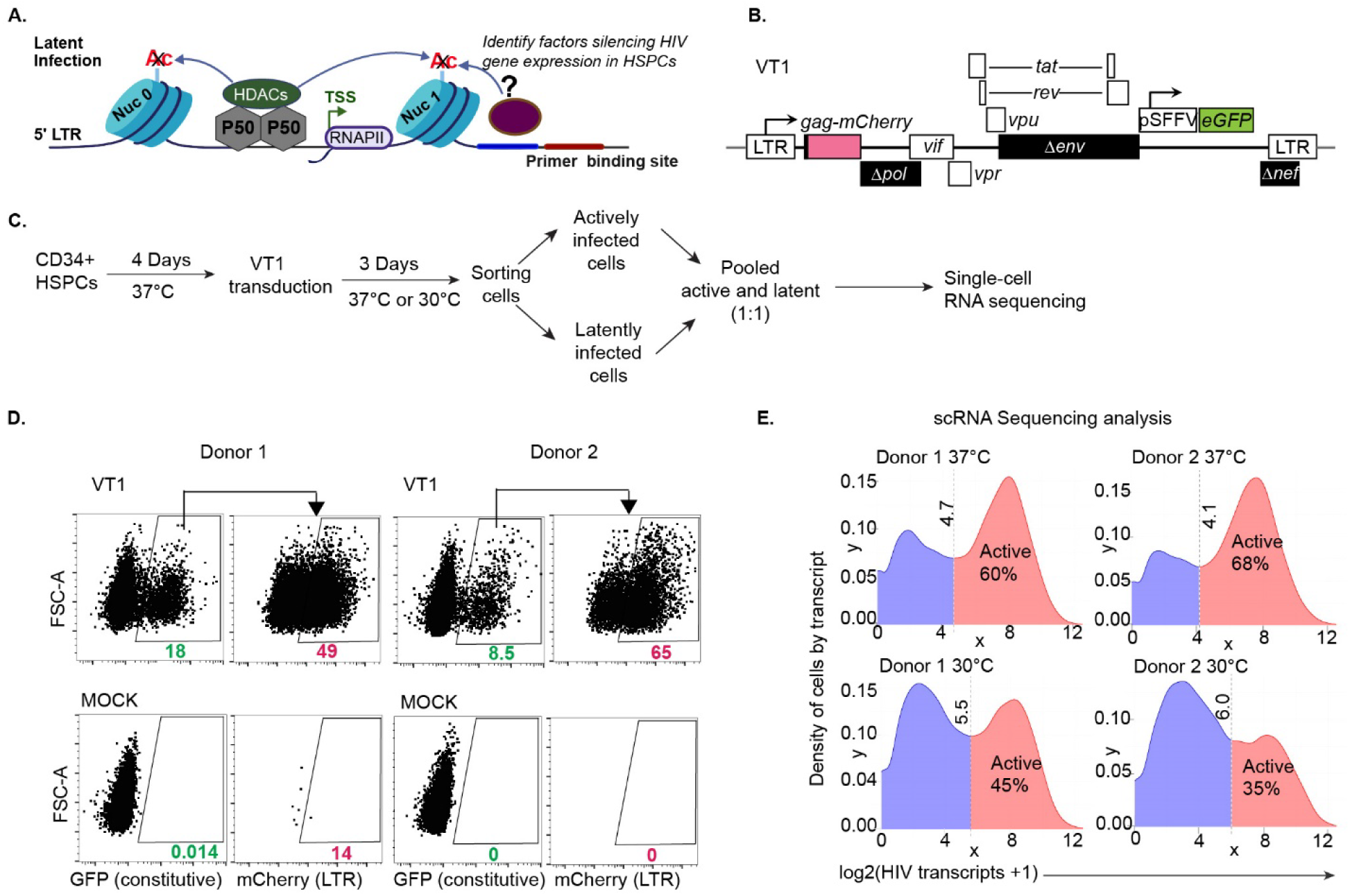
The HIV Reporter VT1 enables identification of latently and actively infected hematopoietic stem and progenitor cells (HSPCs) using flow cytometry and single-cell (sc) RNAseq. **A.** Schematic illustrating the study objective: identification of host factors involved in silencing HIV gene expression in HSPCs. Several histone-modifying enzymes are recruited to the HIV LTR to maintain closed chromatin^102,109^. The primer binding site is depicted as a known target for repressors of retroviral gene expression; nuc, nucleosome; TSS, transcription start site; p50, subunit of NF-kB; HDAC, histone deacetylase; RNAPII, RNA polymerase II; LTR, long terminal repeat. **B.** Schematic diagram of the HIV reporter construct (VT1). Black boxes denote deleted genes; promoter from spleen focus-forming virus (pSFFV); enhanced green fluorescent protein (eGFP). **C.** Experimental workflow illustrating the process used to generate single-cell RNA sequencing datasets. **D.** Flow cytometric analysis of HSPCs transduced with VT1 prior to sorting. Representative plots from Donor 1 and Donor 2 samples at 37°C are shown, illustrating the proportion of transduced cells (GFP+) with LTR activity (mCherry+). The arrow above the flow plots indicates the cell population gated and further analyzed in the subsequent plot. **E.** Density plots of HIV RNA expression from HSPCs treated as diagrammed in C at the indicated culture temperature. Cutoff values (dashed lines) were manually defined near the local minima of each distribution to separate active and latent populations.

Varying culturing conditions lead to different infection outcomes in HSPCs. When cells are grown at 37°C with growth factors including stem cell factor, Flt3 ligand, thrombopoietin, and insulin-like growth factor 1 they actively proliferate and can develop both latent and active HIV-1 infections. In contrast, cells kept at 30°C with the same growth factors remain more quiescent, which promotes the establishment and persistence of latent infections^8,12,14^. To facilitate the identification of host factors that may regulate latency in this cell type, we prepared HSPCs cultured at both temperatures, sorted for infected cells (Fig. 1C, D; Supplementary Fig. S1A), and performed scRNA-seq. To distinguish actively from latently infected cells in the scRNA-seq data, HIV RNA sequences from each cell were log2(x+1) transformed. For each sample, a cutoff threshold was manually selected near the local minimum of the density distribution to separate actively infected cells (higher HIV expression) from latently infected cells (lower expression). Applying this cutoff, the percentage of actively infected cells varied between 30% and 70% among the four samples (Fig. 1E).

### scRNA sequencing reveals differential gene expression between latently and actively infected HSPCs

LIGER (linked inference of genomic experimental relationships)^43,44^ was used to integrate all four single-cell RNA sequencing data sets. This analysis revealed that HSPCs from both donors, cultured at either 37°C or 30°C, were distributed throughout the uniform manifold approximation and projection (UMAP) visualization (Fig. S1B). By analyzing gene expression patterns within each cluster, using canonical blood marker genes such as SPINK2, MPO, LYZ, TCF4, HBD, HDC, RHEX, PF4, and EBF1 (Fig. 2A), we identified the main cell clusters. These clusters represent HSPCs at various stages of hematopoietic differentiation.

**Fig. 2:**
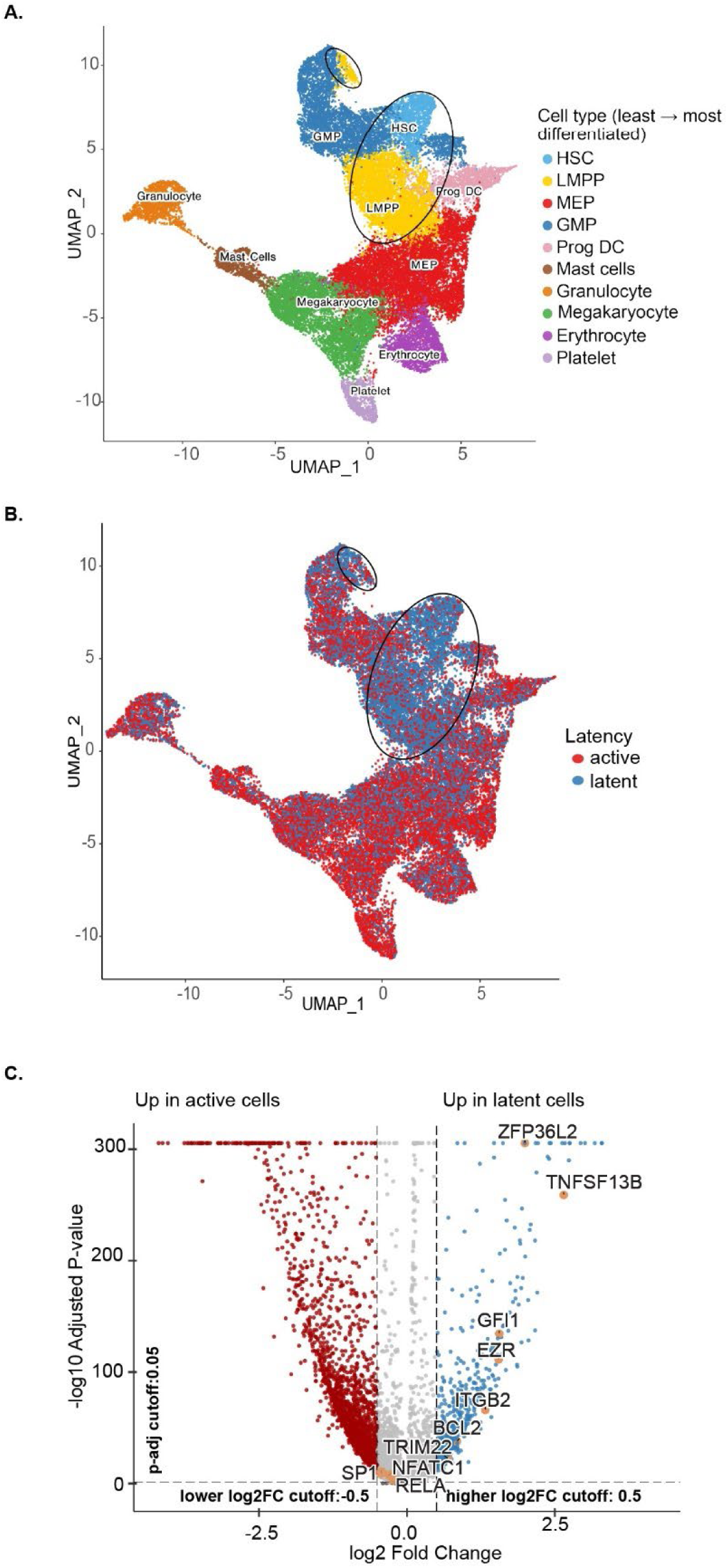
scRNA sequencing reveals differential gene expression between latently and actively infected HSPCs. **A.** UMAP of LIGER-integrated scRNA-seq datasets from the four HSPC samples shown in Fig. 1E. Progenitor cell types were identified based on expression of canonical blood cell marker genes, including *SPINK2*, *MPO*, *LYZ*, *TCF4*, *HBD*, *HDC*, *RHEX*, *PF4*, and *EBF1*. HSC, hematopoietic stem cell; LMPP, lymphoid-primed multipotent progenitor; MEP, megakaryocyte-erythroid progenitor; GMP, granulocyte-macrophage progenitor; prog DC, progenitor dendritic cell. Black ovals highlight cell types that correspond with latent HIV genomes (blue region) in B. **B**. UMAP representation of active and latent HIV-infected cell populations based on the cutoff described in Fig. 1E. **C.** Volcano plot showing genes with differential expression between latently and actively infected cells, as determined by the Wilcoxon rank-sum test. Significant genes were identified using a log2 fold change threshold of 0.5 and an adjusted p-value threshold of 0.05. Genes upregulated in latent infection are shown in blue, while genes upregulated in active infection are shown in red. Genes highlighted as orange dots were selected for further testing.

Cells were classified as actively or latently infected based on HIV expression levels. Specifically, those with log2 HIV expression above the cutoff (Fig. 1E) were designated as actively infected (Fig. 2B). More primitive HSPC clusters, hematopoietic stem cells (HSC) and lympho-myeloid primed progenitor cells (LMPP) predominantly supported latent HIV-1 infection (demarcated region in Fig. 2A, B), while differentiating HSPCs tended to acquire predominantly active infections. Comparative analysis of gene expression profiles between latently and actively infected cells revealed distinct sets of genes upregulated in each condition. Differential expression was assessed using the LIGER runMarkerDEG function and Wilcoxon rank-sum test, with a log2 fold change threshold of 0.5 and an adjusted p-value cutoff of 0.05 (Fig. 2C). Because the role of NF-κB in promoting HIV transcription is well established, we focused on several candidate genes upregulated in HIV latency that are implicated in the regulation of NF-κB signaling; *GFI1, EZR, BCL2, ZFP36L2, TNFSF13B, ITGB2,* and *TRIM22* were selected for further analysis^45–52^.

### GFI1 is an HSPC latency-associated gene that potently silences HIV gene expression in a T cell model of latency

To assess the inhibitory potential of selected genes against HIV expression, the candidate genes were inserted along with an internal ribosome entry site (IRES) into VT1, as illustrated in Fig. 3A. To assess the impact of each cDNA on HIV gene expression, we utilized a T-cell line model of latency, CEM-A2^34,53^. Because we previously established that active infection varies with the virus inoculum^34^, we assessed the proportion of actively infected cells over a range and assessed active versus latent infection flow cytometrically (Fig. 3B-D). Overexpression of GFI1 resulted in a marked reduction in HIV gene expression compared to the BFP control, with inhibition being most prominent at lower amounts of virus (Fig. 3C, D). Additional candidate genes displayed variable inhibition of HIV expression, as summarized by fold changes relative to the control (Fig. 3E and Supplementary Fig. S2). These results indicate that several of the candidate genes can suppress LTR-driven HIV expression to differing extents in CEM-A2 cells.

**Fig. 3:**
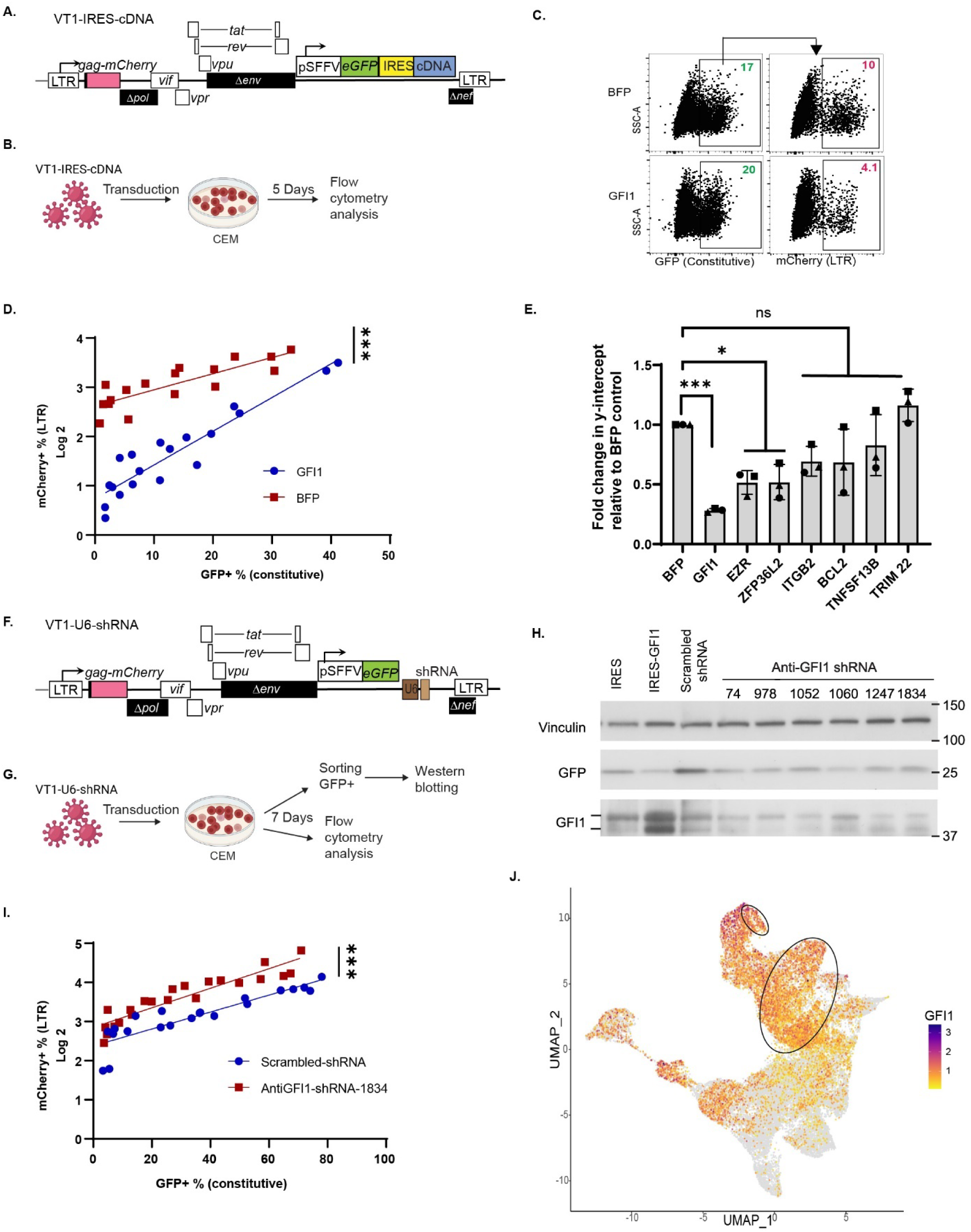
GFI1 is associated with latency in HSPCs and potently silences HIV gene expression in a T cell latency model. **A.** Schematic of the VT1 construct (as in Fig. 1B) with candidate genes (cDNA) inserted after an internal ribosome entry site (IRES) under control of the constitutive (SFFV) promoter. **B.** Experimental outline for analyzing the overexpression of each candidate gene using VT1-IRES-cDNA virus. **C**. Representative flow cytometric plots of LTR-driven and constitutive promoter activity in CEM-A2 cells transduced with the indicated VT1-IRES-cDNA virus. **D**. Summary graph of experiments performed as in C and displayed over a range of virus inoculum (*n*=3 independent experiments). The y-axis shows log2-transformed proportion of cells that had HIV LTR activity (mCherry expression) as a function of the proportion of cells transduced for each inoculum. **E.** Summary graph showing fold changes of y-intercept, comparing BFP control to candidate gene overexpression, as determined by Deming regression (Model II). Data are shown as mean ± standard deviation for *n*= 3 independent experiments. Statistical significance was determined by one-way ANOVA followed by Dunnett’s multiple comparisons test. ***P ≤ 0.001, *P ≤ 0.05. **F**. Schematic of the VT1 construct with an expression cassette containing a U6 promoter inserted upstream of shRNAs as indicated. **G.** Experimental outline for analyzing the silencing of the candidate gene using VT1-U6-shRNA. **H.** Western blot analysis of CEM-A2 cell lysates from cells transduced with viruses containing the indicated GFI1 shRNA (construct in 3F). **I.** Summary graphs as in D except that the indicated shRNA-containing viruses were utilized (*n*=3). **J.** UMAP plot showing GFI1 expression in HSPCs transduced with VT1 from the four integrated scRNA-seq samples described in Figs. 1 and 2.

To confirm this, we tested the hypothesis that knockdown of GFI1 would increase HIV expression. To accomplish this, a short hairpin (sh) RNA expression cassette targeting GFI1 was inserted into the VT1 construct downstream of a U6 promoter (Fig. 3F), and transduced cells were assessed for GFI protein levels via western blot analysis (Fig. 3G, H). One of the shRNAs tested that reduced endogenous GFI1 levels (1834) was further tested and found to increase HIV gene expression, consistent with the conclusion that endogenous GFI1 inhibits viral transcription in these cells (Fig. 3I).

Understanding the interplay between host gene regulation and HIV is critical for elucidating mechanisms of HIV latency. To further investigate how GFI1 levels correlate with HIV activity in HSPCs, we visualized the integrated single-cell RNA-seq datasets and found that GFI1 expression was higher in latently infected HSPC populations (demarcated area shown in Fig. 3J) compared to actively infected cells (see also Fig. 2B). To quantitatively assess this relationship, we performed correlation analysis between host gene expression and HIV gene expression at the single-cell level across the four scRNA-seq samples (Supplementary Fig. S3). GFI1 expression exhibited a negative correlation with HIV gene expression, whereas RELA expression showed a modest positive correlation, further supporting their respective roles as negative and positive regulators of HIV transcription in HSPCs.

### GFI1 targets the HIV sequence adjacent to the primer binding site

GFI1 recognizes a consensus binding sequence, TAAATCAC(A/T)GCA, which is commonly found within the promoters of its target genes^54–56^. To investigate whether the HIV promoter contains a similar GFI1 binding motif, we performed sequence analysis of the HIV LTR. This analysis revealed the presence of a putative GFI1 consensus binding site adjacent to the primer binding site (PBS) (Fig. 4A). Based on this finding, we hypothesized that GFI1 could directly bind to this sequence within the HIV LTR.

**Fig. 4.**
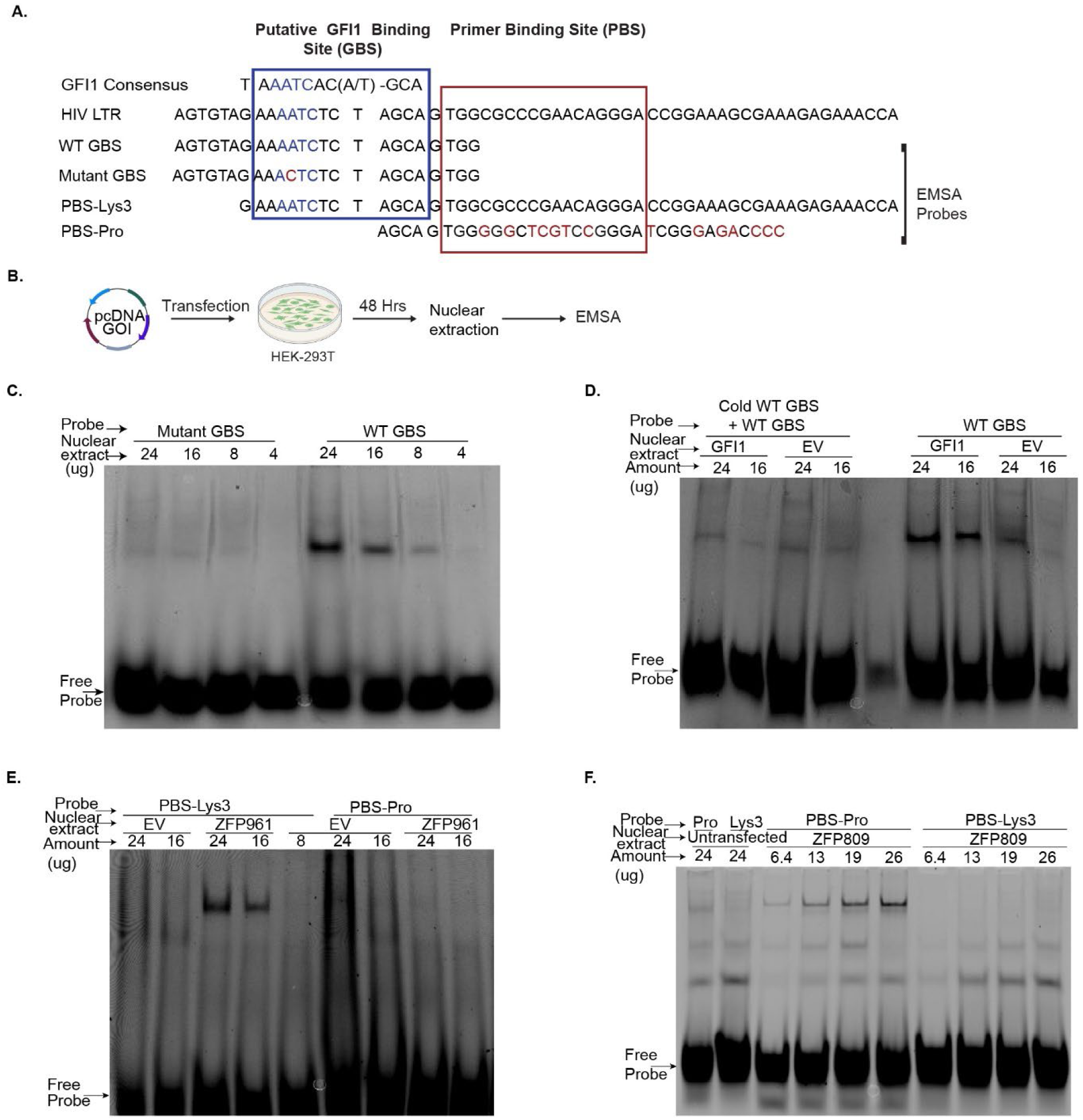
GFI1 binds to a site in the HIV LTR adjacent to the primer binding site (PBS). **A.** Nucleotide sequence of the HIV LTR showing the putative GFI1 binding site (blue rectangle) adjacent to the primer binding site (PBS, red rectangle). The GFI1 binding consensus sequence, Electromobility shift analysis (EMSA) probes for the GFI1 binding site, and individual PBS variants (Lys3 for HIV and Pro for murine leukemia virus (MLV) are also indicated. The mutated GFI1 binding site (GBS) contains an A–to-C substitution previously shown to disrupt GFI1 binding^54^. **B.** Experimental outline for generating nuclear extracts from HEK293T cells**. C**. EMSA of nuclear extracts from HEK293T cells transfected with GFI1-expressing or control plasmid and incubated with the indicated EMSA probe. **D.** EMSA of nuclear extracts prepared as in C, except that an excess of unlabeled (cold) probe was added to the primer incubation step as indicated. **E.** and **F.** EMSA of nuclear extracts prepared as in C except that cells were transfected with ZFP961 (E) or ZFP809 (F) expression plasmids and nuclear extracts were incubated with the indicated PBS EMSA probe. GOI, gene of interest; EV, empty vector

To test this hypothesis, we designed oligonucleotide probes corresponding to the HIV LTR sequence containing the putative GFI1 binding site and performed electrophoretic mobility shift assays (EMSA) (Fig. 4B). Nuclear extracts were prepared from HEK293T cells transfected with pcDNA-GFI1 (overexpressing GFI1) or empty vector (EV). (Fig. 4B). EMSA demonstrated that GFI1 specifically bound to the wild-type LTR-derived probe containing the consensus sequence (Fig. 4C). Consistent with previous reports by Zweidler-McKay et al.^54^, introducing a single A-to-C mutation in the core binding site abolished GFI1 binding to the LTR-derived probe (Fig. 4C). Moreover, competition assays using excess unlabeled wild-type probe further confirmed the specificity of this interaction (Fig. 4D).

Several host factors have been reported to target the PBS of retroviruses, including the mouse Kruppel-associated box (KRAB) zinc finger repressors ZFP961 and ZFP809^24–26^. Numerous studies have shown that accurate pairing between PBS sequences and their respective tRNA molecules facilitates the initiation of reverse transcription^57–61^. However, these PBS sequences can also function as regulatory elements that limit retroviral gene expression. ZFP809, for example, is a murine leukemia virus (MLV) inhibitor that suppresses MLV gene expression by binding to the proline tRNA PBS (PBS-Pro) within the MLV LTR^62^. Similarly, ZFP961, another murine KRAB zinc finger protein, can inhibit Mason-Pfizer monkey virus (which utilizes the tRNA Lys 1,2 PBS [PBS-Lys1,2]) as well as HIV (which uses the Lys3 tRNA PBS [PBS-Lys3] in the HIV LTR) ^26^. As an additional control for our study, we compared the binding specificities of ZFP961 and ZFP809 to the PBS region of the HIV LTR. To do this, we performed EMSAs using probes containing the HIV PBS or a variant containing the PBS-pro, as depicted in Fig. 4A. Consistent with expectations based on prior studies, ZFP961-enriched nuclear extracts bound selectively to the PBS-Lys3 probe, but not the PBS-Pro probe (Fig. 4E). In contrast, ZFP809-enriched nuclear extracts specifically bound the PBS-Pro probe, with no detected binding to PBS-Lys3 sequence found within the HIV LTR (Fig. 4F). These findings confirmed that distinct retroviral PBS region sequences are recognized and targeted by specific host transcriptional repressors, contributing to the selective repression of retroviral transcription and provide a basis for our use of these repressors as controls for subsequent studies of GFI1.

### GFI1 suppresses diverse molecular clones of HIV

HIV is characterized by extensive genetic diversity, with molecular clones exhibiting unique sequence variations that can affect its expression^34^. In addition, variations in HIV sequences can impact interactions with host cellular proteins involved in viral gene expression. To establish a foundation for evaluating the impact of GFI1 across different genetic contexts, we utilized three distinct HIV molecular clones: 89.6 VT1, 454 VT2, and NL4-3 BR3-1 (Fig. 5A). The 89.6 VT1 clone, isolated from the peripheral blood of an individual with AIDS^63^, expresses GFP from constitutive spleen focus-forming virus (SFFV) promoter, and mCherry via control of the HIV LTR (Fig.1B and 5B). In VT1, Nef expression is disrupted by the placement of the SFFV promoter in the *nef* open reading frame (ORF). The 454 VT2 clone^34^ was derived from a person with HIV receiving optimal cART^9^, and in this construct, Nef expression is preserved. Here, the EF1-α constitutive promoter located within the *env* ORF drives mCherry expression (Fig. 5E), and the LTR directs GFP expression. The BR3-1 clone not previously reported and is based on the lab-adapted NL4-3 HIV backbone^64^. BR3-1 features SFFV-driven mCherry expression (inserted in the *nef* ORF) for constitutive transcription and LTR-driven GFP expression (inserted in *env*) as a marker of active infection (Fig. 5H). BR3-1, similar to VT1, does not express Nef. A prior study determined that the presence or absence of a *nef* open reading frame did not affect the proportion of latent or active infection^34^. Utilizing reporter constructs derived from three HIV molecular clones provides a robust framework for comparing the effects of GFI1 on HIV across naturally occurring sequence variations.

**Fig. 5.**
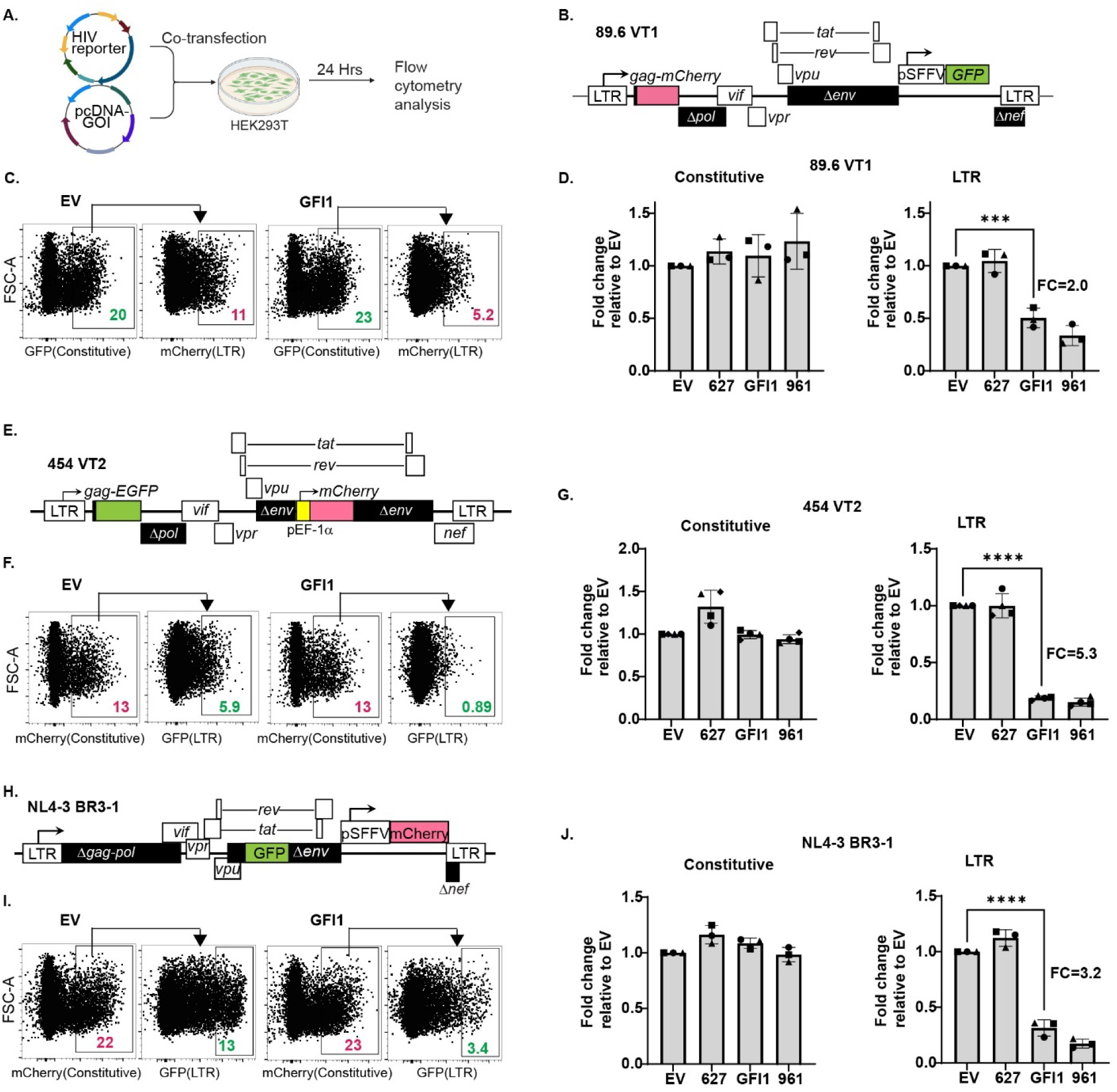
GFI1 suppresses diverse molecular clones of HIV. **A**. Experimental workflow for testing the effect of GFI1 on different HIV clones in HEK293T cells**. B.** Schematic of the 89.6 VT1 construct (as in Fig. 1B). **C.** Representative flow cytometric plots of transfected HEK293T cells. Active LTR expression was determined by quantifying the proportion of GFP-expressing cells that were mCherry-positive. **D.** Summary graphs of flow cytometry data represented in C. **E.** Schematic of the 454 VT2 construct in which the LTR drives expression of a Gag-GFP fusion protein. The constitutive promoter pEF-1α drives mCherry inserted in the *env* reading frame. **F.** Representative flow cytometric plots of transfected HEK293T cells. Active expression was determined by quantifying the proportion of mCherry-expressing cells that were GFP-positive. **G.** Summary graphs of flow cytometry data represented in F**. H.** Schematic of the NL4-3 BR3-1 HIV construct. LTR drives GFP inserted in the *env* reading frame. Constitutive promoter pSFFV drives mCherry inserted in the *env* and *nef* reading frames. **I.** Representative flow cytometric plots of transfected HEK293T cells. Active expression was determined by quantifying the proportion of mCherry-expressing cells that were GFP-positive. **J.** Summary graphs of flow cytometry analysis represented in I. For D, G and J, fold change was calculated by dividing the expression observed with the test gene by the expression observed with the EV control. Data are shown as mean ± standard deviation for *n*=3 independent experiments in panels D and J, and *n =* 4 in panel G, performed in triplicate. Statistical significance was determined by one-way ANOVA followed by Dunnett’s multiple comparisons test. ***P ≤ 0.001 and ****P ≤ 0.0001. See Supplementary Fig. S4 for additional flow cytometry plots corresponding to panels C, F, and I. GOI, gene of interest; EV, empty vector.

To investigate the impact of GFI1 on genetically distinct HIV reporter constructs, we co-transfected each with either pcDNA-EV, GFI1, positive control KRAB zinc finger protein ZFP961, or a related KRAB zinc finger protein that serves as a negative control (ZNF627). As expected, none of these factors reduced constitutive promoter-driven expression of VT1 compared to the EV control (Fig. 5C, D). In contrast, both GFI1 and ZFP961 markedly suppressed LTR-driven transcription in VT1. Consistently, similar inhibition of LTR activity by GFI1 and ZFP961 was observed in VT2 and BR3-1 clones, with negligible impact on constitutive promoter activity (Fig. 5F, G, I, J, and Supplementary Fig. S4). As expected, ZNF627 did not exert any inhibitory effect on LTR-mediated activity in any of the clones tested. These findings demonstrate that GFI1 specifically inhibits HIV LTR-driven expression, without affecting constitutive promoter activity, underscoring its regulatory role across genetically diverse HIV clones.

### GFI1 silences HIV gene expression through its DNA-binding and corepressor-interaction domains

GFI1 contains three distinct domains that mediate interactions with various transcriptional regulators. The N-terminal SNAG domain recruits corepressors such as REST corepressor 1 (CoREST), lysine-specific demethylase 1 (LSD1), and histone deacetylases (HDACs), which together form the CoREST complex^65–68^. The central domain interacts with additional factors, including G9a and additional HDACs^69,70^, and is also known to associate with NF-κB^45^.

The C-terminal domain of GFI1 contains six C2H2-type zinc fingers, which facilitate binding to target genes. Among these, fingers 3–5 are required for recognition and binding of its target DNA sequences^54,71,72^ (Fig. 6A). To assess the necessity of the DNA-binding domain for GFI1-mediated suppression of HIV LTR activity, we introduced the N382S point mutation into the fifth zinc finger, a mutation previously shown to disrupt DNA binding^73–75^. Western blot analysis confirmed that GFI1 protein expression was not reduced by the N382S mutation (Fig. 6B). To evaluate the functional consequences of the mutation, we co-transfected either wild-type GFI1 or the N382S mutant together with VT2 in HEK293T cells and measured LTR activity. Compared to wild-type GFI1, the N382S mutant displayed a significant reduction in its ability to suppress LTR-driven transcription (Figs. 6C, D).

**Fig. 6.**
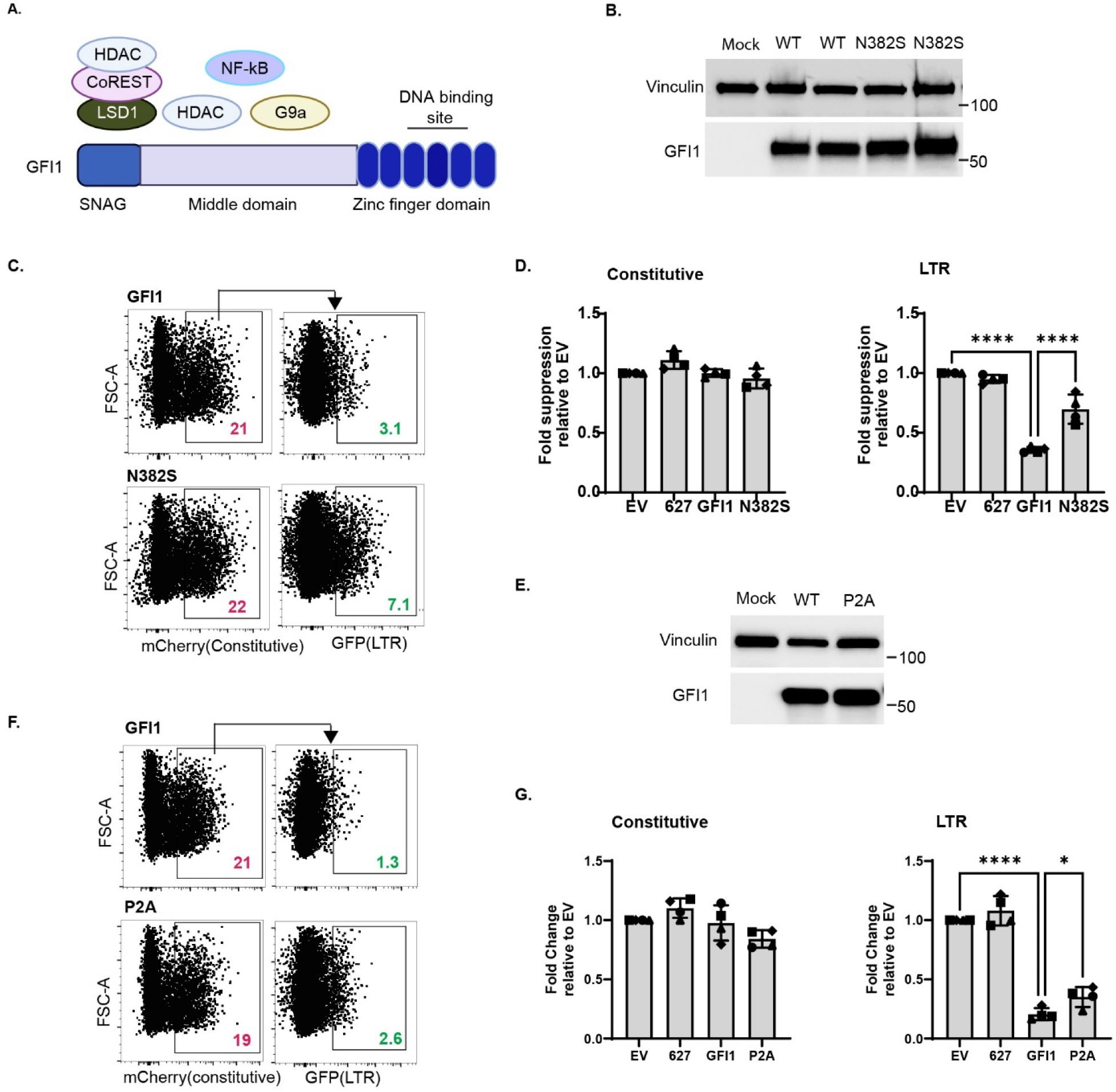
Functional GFI1 zinc finger and SNAG domains are required for maximal repression of HIV transcription. **A.** Diagrammatic illustration of the domains of GFI1 with associated regulatory proteins. CoREST, REST corepressor 1^66^; HDAC, histone deacetylase^69,70^; LSD1, lysine-specific demethylase 1^65,66^; G9a, euchromatic histone methyltransferase 2 (EHMT2)^69^; NF-κB, nuclear factor kappa-light-chain-enhancer of activated B cells^45,110^. **B.** Western blot analysis of HEK293T cells transfected with the indicated wild-type or mutant GFI1 construct. The N382S mutation disrupts GFI1 binding to DNA^73,75^. **C.** Representative flow cytometric plots of HEK293T cells co-transfected as in Fig. 5A. The N382S mutant or wild-type GFI1 expression plasmid was utilized. **D.** Summary graphs of flow cytometric data shown in C. **E.** Western blot analysis of HEK293T cells transfected as in part B with the indicated wild-type or mutant GFI1 construct. The P2A mutation disrupts the SNAG domain of GFI1 shown in part A and inhibits binding of LSD1^66,67^. **F.** Representative flow cytometric plots of HEK293T cells co-transfected as in Fig. 5A with wild-type or mutant GFI1 constructs. **G.** Summary graphs as in D. For D and G, fold change was calculated by dividing the expression observed with the test genes by the expression observed with the EV. Data are presented as mean ± standard deviation for *n*=4 independent experiments performed in triplicate. Statistical significance was determined by a one-way ANOVA followed by Sidak’s multiple comparisons test. *P ≤ 0.05; ***P ≤ 0.001; and ****P ≤ 0.0001.

Because GFI1 utilizes its SNAG domain to recruit corepressors, including LSD1 and HDACs to target gene sites and enable transcriptional repression^66,67,70^, we hypothesized that the SNAG domain is important for GFI1-mediated suppression of the HIV LTR. To test this, we introduced a P2A mutation that disrupts this domain^66,67^. Western blot analysis confirmed that this P2A mutation did not reduce GFI1 expression levels in HEK293T cells (Fig. 6E). To determine whether the P2A mutation alters GFI1 activity, we co-transfected VT2 in HEK293T cells with either wild-type GFI1 or the P2A mutant and measured LTR activity. LTR-driven transcription was considerably less inhibited by the P2A mutant relative to wild-type GFI1, indicating that an intact SNAG domain is important for maximal suppression of HIV LTR activity by GFI1 (Figs. 6F, G). Notably, the P2A mutant still exhibited some degree of suppression compared to the empty vector, indicating that GFI1 utilizes its internal co-repressor interaction domains to suppress HIV transcription (Fig. 6A)

### GFI1 antagonizes HIV-activating transcription factors Tat and NF-kB

The viral Tat protein serves as a master regulator of HIV gene expression, functioning as a transactivator (Fig. 7A). Tat binds to the transactivation response (TAR) element in nascent HIV transcripts and recruits the positive transcription elongation factor b (P-TEFb) complex, thereby enhancing the efficiency of viral RNA elongation by RNA polymerase II ^31,32,76–78^. Additionally, Tat interacts with nuclear histone acetyltransferases (HATs), resulting in increased acetylation of histones and alterations in chromatin structure around the HIV long terminal repeat (LTR), further facilitating viral transcription^79–81^. In contrast, the cellular protein GFI1 acts as a transcriptional repressor by recruiting HDACs, a mechanism that may also occur at the LTR region, leading to deacetylation of histones, a condensed chromatin structure, and limited viral gene expression. Importantly, GFI1 is itself subject to acetylation, and this modification can inhibit its ability to interact with binding partners^82^. Thus, increased acetylation driven by Tat-associated HATs may diminish GFI1’s repressive function.

**Fig. 7.**
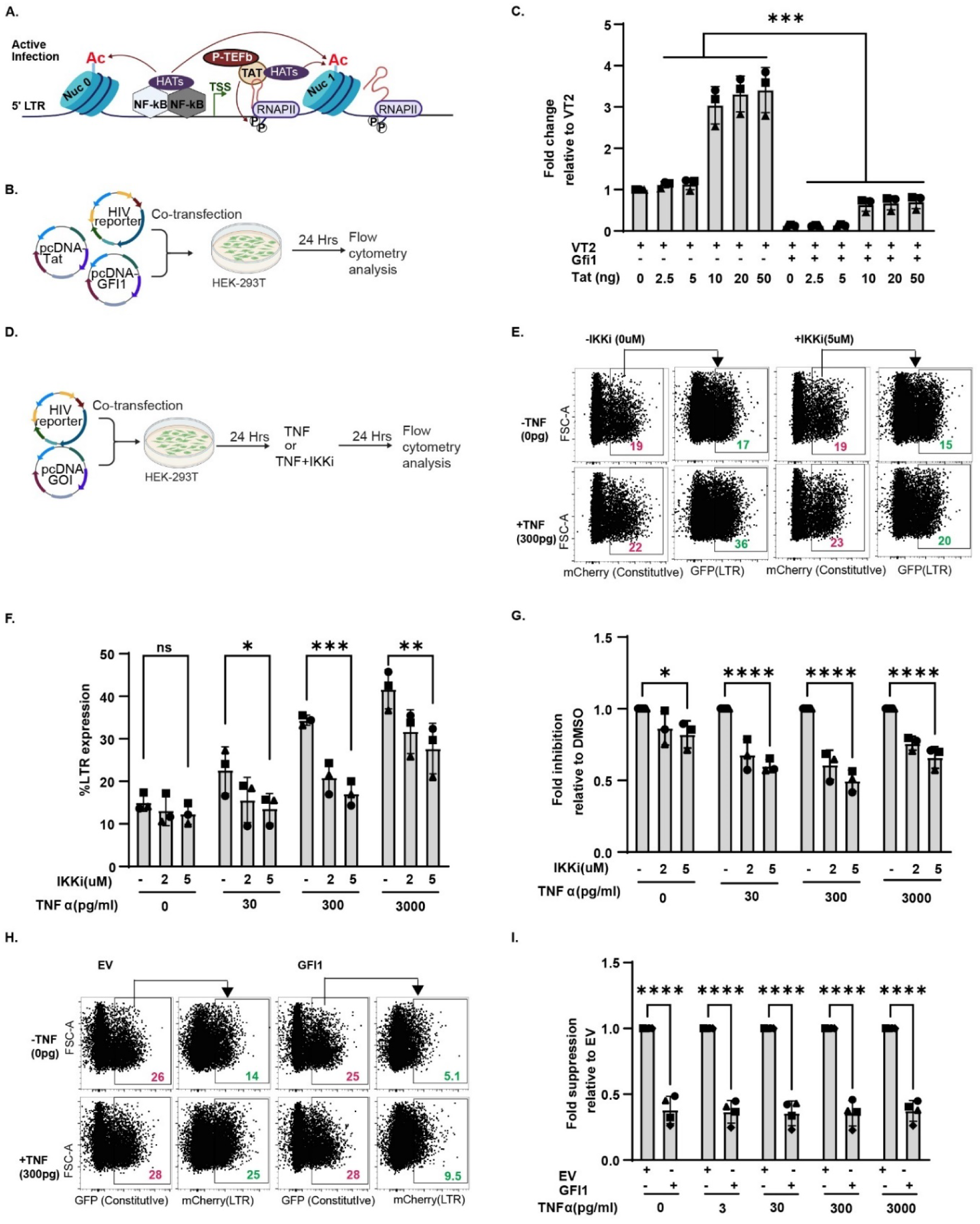
GFI1 antagonizes Tat and NF-kB-dependent activation of the HIV LTR. **A.** Diagrammatic representation of the HIV transactivator of transcription (Tat) and nuclear factor kappa-light-chain-enhancer of activated B cells (NF-κB) activation of HIV gene expression at the LTR^31,32,76,102,103^. Ac, acetylation; HATs, histone acetyltransferases; P-TEFb, positive transcription elongation factor b; TAR, trans-activation response element; RNAPII, RNA polymerase II; circled P, phosphorylation. **B.** Experimental workflow for testing the effect of GFI1 on Tat-mediated activation of VT2. **C.** Summary graph of flow cytometric data of VT2 LTR activity in HEK293T cells transfected as shown in B with the indicated amount of Tat or GFI1. Fold change is LTR-driven expression of each sample normalized to that of VT2 alone. See Supplementary Fig. S5B for summary graph of constitutive promoter activity. **D.** Experimental workflow for testing the effect of GFI1 on NF-κB–mediated activation of VT2. **E.** Representative flow cytometric plots of HEK293T cells transfected as in D with the addition of IkB kinase inhibitor (IKKi), and tumor necrosis factor a (TNF-α) as indicated. **F**. Summary graph of flow cytometry data shown in E using HEK293T cells treated as indicated and as shown in D. **G.** Summary graph of data from E as fold change relative to the untreated sample. See also Supplementary Fig. S5C for constitutive promoter activity. **H.** Representative flow cytometric plots of HEK293T co-transfected with VT2 treated as shown in D plus GFI1or EV as indicated. **I.** Summary graph of flow cytometry data shown in H. See also Supplementary Fig. S5D for constitutive promoter activity. Data are shown as mean ± standard deviation for *n*=3 independent experiments performed in triplicate. Statistical significance was determined by a two-way ANOVA (C, F, G, and I) followed by Sidak’s multiple comparisons test. **P ≤ 0.01; and ****P ≤ 0.0001. GOI, gene of interest; EV, empty vector.

Based on these prior findings, we hypothesized that there may be antagonistic interplay between these viral and host regulatory proteins in modulating HIV transcription. To test this hypothesis, we assessed whether GFI1 could suppress Tat-induced activation of the HIV LTR by co-transfecting HEK293T cells with VT2 and GFI1, alongside increasing amounts of Tat expression plasmid (Fig. 7B). Our results show that GFI1 expression significantly reduced Tat-induced HIV LTR activity compared to cells receiving reporter construct alone, indicating that GFI1 is able to counteract Tat-mediated transactivation of the viral promoter (Fig. 7C). Consistent with our hypothesis of an antagonistic interplay, at high enough levels of Tat, the repressive effect of GFI1 was diminished. Neither Tat nor GFI1 similarly altered expression of the constitutive promoter (Supplementary Fig. S5A, B).

NF-κB is a central regulator of HIV transcription, exerting its effects by binding to specific sites within the HIV LTR promoter. One of the mechanisms underlying NF-κB–mediated activation of the LTR is the recruitment of histone acetyltransferases (HATs), which acetylate histone tails and thereby regulate chromatin structure to facilitate transcriptional activation^83–85^. It is possible that the transcriptional repressor GFI1 could counter the activity of NF-κB by bringing histone deacetylases (HDACs) to the promoter, thereby potentially competing with the chromatin-modifying effects of NF-κB. Alternatively or in addition, GFI1 can act as a negative regulator of NF-κB by physically binding NF-κB and disrupting its ability to bind its target sites^45^. Furthermore, studies indicate that GFI1 can contribute to the inhibition of NF-κB activation in inflammatory conditions^46^.

To this end, we explored whether GFI1 inhibits NF-κB–mediated HIV activation. We first assessed the effects of NF-κB activation and inhibition on VT2 LTR-driven expression. To stimulate NF-κB, we treated HEK293T cells transfected with the reporter construct with TNF-α. To block NF-κB activation, we used an IκB kinase (IKK) inhibitor (IKKi)^86^ (Fig. 7D). IKK is a key component of the NF-κB signaling pathway, responsible for phosphorylating IκB proteins and thereby enabling NF-κB activation^87,88^. Inhibiting IKK allows us to test the impact of reduced NF-κB activity on HIV LTR-driven expression. Together, these treatments allowed us to assess how NF-κB regulates VT2 LTR activity in the presence or absence of GFI1. (Fig. 7D). TNF-α stimulation led to robust activation of LTR-driven expression, as measured by VT2 LTR activity (Fig. 7E, F). When IKKi was added, we observed a dose-dependent reduction in VT2 activity (Fig. 7F, G). As expected, TNF and IKKi did not alter expression from the constitutive promoter (Supplementary Fig. S5C).

Next, we investigated whether GFI1 could similarly suppress NF-κB–driven LTR activation (Fig. 7D). Co-expression of GFI1 reduced VT2 expression both in the presence and absence of TNF-α, demonstrating inhibitory effects akin to those observed with IKKi on TNF-α–induced activation, as well as suppression under basal conditions (Fig. 7H and 7I and Supplementary Fig. S5D). These findings indicate that GFI1 can antagonize NF-κB-dependent LTR-driven HIV transcription.

### A combination of histone deacetylase inhibition and activation of NF-κB-dependent signaling pathways effectively reverses GFI1-mediated repression of HIV gene expression

Given that GFI1 interacts with corepressors such as HDACs, LSD1, and G9a^66,69,70,89,90^, and antagonizes NF-κB signaling^45,46^, we hypothesized that inhibition of these GFI1-associated factors would reverse GFI1-mediated suppression of LTR-driven expression. Notably, GFI1 is known to associate with a variety of different class I histone deacetylases, including HDAC1, which plays a key role in transcriptional repression of HIV-1 ^66,69,70,91^. To target these enzymes, we utilized entinostat, a selective inhibitor of class I HDACs^92,93^; GSK-LSD1, which interferes with the demethylase activity of LSD1 ^90,94^; A-366, an inhibitor of G9a methyltransferase activity^95,96^, and bryostatin 1 protein kinase C agonist that activates NFκB ^97,98^.

Accordingly, CEM-A2 cells transduced with increasing amounts of VT1 virus expressing either GFI1 or a BFP control (Fig. 3A) were treated with these LRAs as depicted in Fig. 8A. Notably, the combination of all the LRAs induced the greatest response, resulting in a 35-fold increase in reporter activity in GFI1-expressing cells versus a 13-fold increase in BFP controls (Fig. 8B, C; Supplementary Table S1). Among individual agents, the HDAC inhibitor entinostat produced the strongest reversal of GFI1-induced repression (summarized in Fig. 8C and Supplementary Table S1), yielding a 13.8-fold increase in LTR reporter activity for VT1-IRES-GFI1-expressing cells as measured by changes in the y-intercept across a range of virus inoculum compared to solvent control. This compares to 6.3-fold for the VT1-IRES-BFP control. Bryostatin alone had a modest effect, increasing expression by 3.2-fold with GFI1 and 2.2-fold with BFP. The impact of inhibiting demethylase activity via LSD1 inhibitor and methyltransferase activity via A366 was small and only statistically significant in combination with entinostat or bryostatin (Fig. 8C). Across treatments, the magnitude of reversal was consistently more pronounced in GFI1-expressing cells than in BFP controls, indicating that targeting corepressors and signaling pathways influenced by GFI1 effectively diminishes its inhibition of HIV LTR activity.

**Fig. 8.**
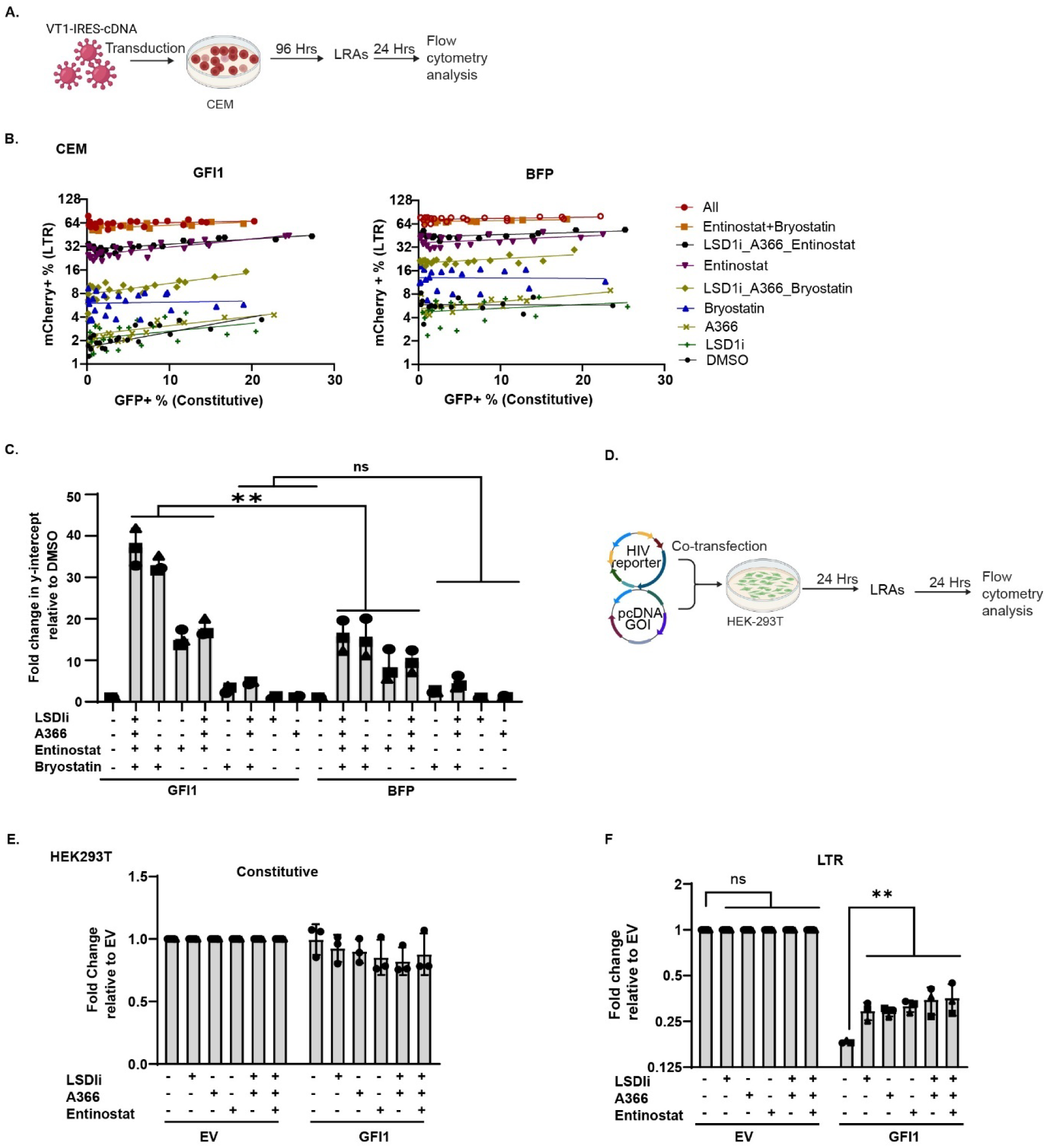
Small molecules that alter histone acetylation and NF-κB activation reverse GFI1-mediated repression of HIV gene expression. **A.** Experimental workflow for testing the effect of latency reversal agents (LRAs) on GFI1-mediated inhibition of LTR-driven expression in CEM-A2 cells. **B.** Summary graphs of CEM-A2 cells transduced with the indicated virus containing GFI1 or control (BFP) and treated with the indicated LRAs. Individual data points are shown from 3 independent virus titrations and LRA treatment experiments. **C.** Summary graph of fold changes in y-intercept, obtained from Deming regression of log2-transformed virus titration data shown in panel B. See also Supplementary Table S1 for comprehensive results. **D.** Experimental workflow for testing the effect of latency reversal agents (LRAs) on GFI1-mediated inhibition of LTR-driven expression in HEK293T cells. **E.** and **F.** Summary graphs of flow cytometric data for the constitutive (E) and LTR-driven (F) fluorophores. Data is shown as mean ± standard deviation for *n*=3 independent experiments performed in triplicate. Statistical significance was determined by a one-way ANOVA followed by Sidak’s multiple comparisons test. ****P ≤ 0.0001, **P ≤ 0.01. GOI, gene of interest; EV, empty vector.

To determine the effects of the LRAs on GFI1-mediated repression across cell types, HEK293T cells were co-transfected with the VT2 reporter and either GFI1 or empty vector control, followed by treatment with the indicated LRAs (Fig. 8D). The LRAs diminished GFI1-mediated suppression of LTR-driven reporter expression. Unlike in CEM-A2 cells, where certain LRAs were much more effective than others, all LRAs tested in HEK293T cells similarly diminished GFI1-mediated suppression of LTR-driven reporter expression (Fig. 8E, F; Supplementary Fig. S6B). Thus, the cell type of origin and/or the state of the viral LTR (plasmid or integrated provirus) influences the extent to which cellular factors can regulate the HIV LTR.

## DISCUSSION

Although accumulating evidence indicates that HSPCs serve as a critical HIV reservoir, the mechanisms underlying latency within these cells remain largely unresolved^7,9,99^. To expand our understanding of HIV latency in HSPCs, we utilized single-cell RNA sequencing to comprehensively profile host factors associated with HIV latency in these cells. We applied the 89.6 VT1 HIV reporter system to distinguish between latently and actively infected HSPCs on the basis of RNA expression^34^. Our single-cell transcriptomic analyses revealed a subset of genes increased in HIV latency. From this subset, we functionally examined genes previously implicated in the regulation of NF-κB signaling, including GFI1, EZR, ZFP36L2, ITGB2, BCL2, TNFSF13B, and TRIM22^45–52^. Among these, GFI1 exhibited the strongest suppressive effect on HIV gene expression in the CEM T cell latency model. Interestingly, the finding that GFI1 RNA expression correlated better with latency in HSPCs than NF-κB suggests that gene silencing mediated by repressors may be more important than NF-kB activity for determining latency establishment in this cell type. Accordingly, our investigation focused on the mechanisms by which GFI1 mediates silencing of HIV transcription, although further research will be required to elucidate the roles of the other identified genes.

GFI1 is a critical transcription factor in hematopoiesis, regulating the cell fate decisions of hematopoietic stem cells in the lymphoid and myeloid lineages^100^. GFI1 is highly expressed in HSCs but is lost upon differentiation, which may help explain why HIV establishes latency in HSCs. As a DNA-binding protein, GFI1 employs a C2H2-type, C-terminal zinc finger domain to recognize consensus sequences within its target DNA sequences. Notably, we identified a GFI1 binding site within the HIV 5’ LTR^54,72^, adjacent to the PBS, suggesting a role of GFI1 in regulating HIV latency at the proviral level. We demonstrated that GFI1 directly engaged this HIV LTR sequence, establishing a mechanistic framework by which GFI1 orchestrates transcriptional silencing. This supports the concept that this region around the PBS functions as a regulatory hub for host-mediated retroviral silencing. The regulatory importance of the PBS region in HIV silencing is further supported by our findings and previous reports demonstrating that transcriptional repressor mouse ZFP961 can silence HIV via the PBS^25,26^. Additionally, mouse ZFP809 as well as human ZNF417 and ZNF587 have been reported to repress retroviral gene expression through PBS-dependent mechanisms^24–26,62,101^, thereby reinforcing the importance of this locus in maintaining viral latency. Taken together, our findings highlight the PBS region as a central hub for host factor-mediated regulation and suggest that variations in this region may influence both host susceptibility and retroviral latency. To further investigate the mechanism underlying GFI1-mediated repression of HIV transcription, we examined the role of its DNA-binding domain. Point mutation analysis revealed that an N382S mutation in the fifth zinc finger, which disrupts GFI1’s recognition of target DNA sequences, relieves suppression of HIV transcription, despite normal protein expression levels. This confirmed that DNA binding is important for GFI1 to exert its repressive effects, a result consistent with prior studies mapping target gene recognition to zinc fingers 3 to 5 ^54,71,72^. Thus, without a functional DNA-binding interface, GFI1 is less able to silence proviral transcription, highlighting the centrality of sequence-specific DNA recognition in its role as a transcriptional repressor.

The integrity of GFI1’s N-terminal SNAG domain, which recruits chromatin-modifying co-repressors such as LSD1 and HDACs^66,67,70^ is also necessary for efficient transcriptional repression. Introduction of the P2A mutation, which disrupts SNAG-mediated interactions, resulted in significant derepression of HIV LTR transcription relative to wild-type GFI1. These data establish that maximal GFI1-mediated repressive chromatin assembly is dependent on an intact SNAG domain, reinforcing its function as a molecular scaffold for the CoREST complex and related chromatin-modifying enzymes ^66,69,89^. This requirement for co-repressor recruitment via SNAG complements the necessity of DNA-binding, suggesting that efficient proviral silencing by GFI1 relies on both sequence-specific localization and the subsequent assembly of epigenetic silencers at the HIV LTR. Residual GFI1 activity present despite mutation of its SNAG highlights the importance of additional GFI1 domains capable of interacting with corepressors, as well as its capacity to inhibit NF-κB as discussed below.

Because GFI1 mediates chromatin silencing, while factors such as NF-κB and Tat promote transcriptional activation partly through chromatin remodeling, we investigated how these opposing forces interact at the HIV LTR^79,80,102,103^. GFI1 markedly reduced Tat-driven LTR activity, though higher Tat levels could overcome this effect, revealing a threshold in the balance between activation and silencing. Similarly, GFI1 suppressed NF-κB–dependent transcription under both basal and TNF-α–stimulated conditions, an inhibition comparable to the effects of pharmacological NF-κB blockade. These findings identify GFI1 as a key regulator of HIV latency that can counteract major viral and host transcriptional activators. Because prior studies have reported direct interactions between NF-κB and GFI1^45^, further investigation is warranted to determine whether such interplay contributes to GFI1-mediated repression of HIV gene expression.

Efforts to reverse HIV latency must address not only the epigenetic barriers imposed by host factors like GFI1, but also the interplay with transcriptional activators such as NF-κB. Our findings reinforce that effective latency reversal is not achieved through counteracting the repression mediated by repressors alone, but also on robust stimulation of transcriptional activators. For example, bryostatin, a well-known LRA that acts as a protein kinase C (PKC) agonist, thus activating NF-κB-driven transcription from the HIV LTR^97,104,105^, counteracted the GFI1-mediated transcriptional repression. Treatment with the class I HDAC inhibitor entinostat effectively reversed GFI1-mediated repression compared to other LRAs alone, suggesting that in CEM cells, GFI1 mainly recruits HDACs to the HIV LTR to silence transcription. The limited efficacy of LSD1 demethylase inhibition observed in 293T cells, but not CEM T cells, is consistent with recent results showing that LSD1’s role as a demethylase enzyme is not strictly necessary for GFI1 to function as a transcriptional repressor. LSD1’s role may instead be to serve as a scaffold protein, facilitating the attachment of enzymes like HDACs to GFI1^106,107^. Inhibition of the methyltransferase G9a similarly demonstrated cell type-specific effects indicating that G9a may not be involved in maintaining latency in T cell systems. The differential responsiveness observed across cell types points to the influence of cellular environment and chromatin context on the effectiveness of these interventions.

Our observations indicate that latency reversal agents (LRAs) are most effective when employed in combination, underscoring the multifactorial nature of HIV latency and the necessity of simultaneously targeting both repression and activation mechanisms. Notably, the combination of bryostatin and entinostat stood out for its ability to overcome deficiencies in GFI1-mediated transcriptional regulation, suggesting a strong synergistic effect between these agents. This finding is consistent with our prior study utilizing these LRAs in HSPCs^13^ as well as studies in other latency models^28,108^. The magnitude of LRA-mediated reversal was substantially greater in GFI1-expressing cells compared to controls, affirming that these agents target specific mechanisms associated with GFI1-driven repression, as illustrated in the working model in Fig. 9. Collectively, these insights support a treatment strategy that combines activation of host transcriptional pathways like NF-κB with targeted disruption of chromatin-based repression for robust and context-dependent reactivation of latent HIV.

**Fig. 9:**
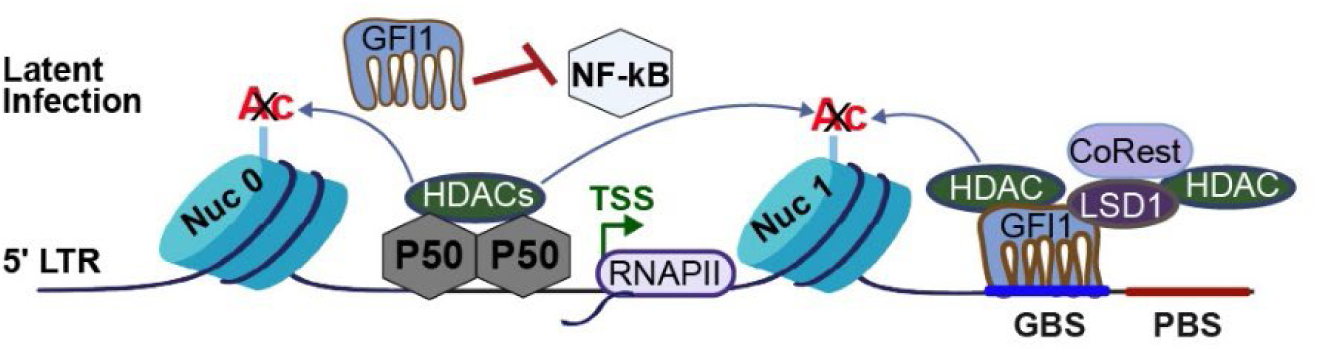
Working model of GFI1 silencing of HIV gene. GFI1 binds to the HIV LTR and recruits histone modifiers that promote closed chromatin and inhibit NF-κB activity, thereby silencing HIV gene expression. Nuc, nucleosome; p50, NF-κB p50 subunit; HDACs, histone deacetylases; TSS, transcription start site; RNAPII, RNA polymerase II; GBS, GFI1 binding site; PBS, primer binding site; CoREST, corepressor for REST; LSD1, lysine-specific demethylase 1.

Prior findings by Painter et al.^12^ showed that quiescent HSPCs, maintained in their undifferentiated state at lower temperature, are more likely to retain latent HIV infection. In our study, we observed that the HSC cluster, characterized by fewer markers of differentiation as identified by scRNA-seq, exhibited a high prevalence of latent HIV infection, whereas HSPC clusters with markers indicating commitment toward specific differentiation lineages favored active viral replication. These results support a model in which HSPCs residing in the bone marrow niche, characterized by cellular quiescence and lower temperatures, may represent an important reservoir for latent HIV. This differentiation-dependent dichotomy suggests that the HIV reservoir is dynamically shaped by stem cell hierarchies, raising essential considerations for cure strategies aimed at purging latent reservoirs.

Moreover, selective expression of GFI1 RNA in HSC clusters supports our finding that GFI1 serves an important role in latency establishment. Finally, the evolutionary conservation of the GFI1 binding site in the HIV genome suggests the intriguing possibility that HIV-1 benefits by maintaining latency in HSCs. Indeed, prior work from our group demonstrated that HIV can be spread as a latent genome to differentiated cell types via cellular proliferation^9,15^, providing HIV with an alternative way of replicating its genome that protects it from immune recognition and viral cytopathic effects.

In addition to GFI1, our single-cell RNA sequencing approach uncovered several other candidate genes that could be associated with HIV latency, opening avenues for further exploration of alternative silencing pathways. It is worth noting, however, that upregulation of candidate genes in latency may reflect both upstream regulation of latency and suppression downstream of HIV gene expression in actively infected cells. Additional functional studies will be required to distinguish these possibilities.

One limitation of our mechanistic analysis of GFI1-mediated transcriptional repression is that our conclusions are primarily drawn from cell line models, which may not fully recapitulate the nuanced biology or the complexity of HIV latency observed in vivo. Additionally, variability in responses to LRAs we observed across different cellular contexts (HEK293T cells versus CEM cells) indicates that strategies effective in one cell type may not generalize broadly. A deeper understanding of how latency varies across cell differentiation hierarchies will be essential to advancing approaches for reservoir eradication.

Our work illuminates the dynamic balance of silencing and activating forces at the HIV LTR, with GFI1-mediated repression antagonizing activation by host and viral transcriptional activators. The region around the PBS emerges as a regulatory hub, where host factor interactions decisively shape transcriptional outcomes. The ability of LRAs to overcome GFI1-mediated repression, particularly through targeted inhibition of HDAC recruitment and activation of NF-κB hint at actionable pathways for latency reversal targeting GFI1. Continued research focused on the regulatory circuitry at the HIV LTR, the contribution of additional silencing candidates, and confirmatory mechanistic studies in relevant cellular reservoirs will be pivotal. These steps are essential prerequisites for translating findings to in vivo models and refining cure strategies, whether aimed at durable silencing (“block and lock”) or reactivation and clearance (“shock and kill”). Collectively, these insights move the field closer to tailored interventions for the effective targeting of the latent HIV reservoir.

## METHODS

### Ethics Statement

Anonymized cord blood cells were obtained from New York Blood Center after obtaining informed consent. Studies using these cells were determined to be exempt from human studies requirements by the University of Michigan Institutional Review Board because the project involves only biological specimens that cannot be linked to a specific individual by the investigator(s) directly or indirectly through a coding system.

### Cell culture

All cells were maintained at 37°C in a humidified atmosphere with 5% CO₂ unless otherwise indicated. HEK293T (ATCC CRL-3216) cells were cultured in D10 medium composed of Dulbecco’s Modified Eagle Medium (DMEM, Gibco, cat# 11995073) supplemented with 100 units/mL penicillin, 100 µg/mL streptomycin, 2 mM glutamine (Pen-Strep-Glutamine, Invitrogen, cat# 10378016), 10% fetal bovine serum (FBS, Gibco, cat# 26140079), and 2.5 µg/mL plasmocin (InvivoGen, cat# ant-mpp) to prevent mycoplasma contamination. CEM-A2 cells^53^ were cultured in R10 medium (RPMI-1640, Gibco, cat# 11875093) supplemented with 100 units/mL penicillin, 100 µg/mL streptomycin, 2 mM glutamine, 10% FBS, and 2.5 µg/mL plasmocin.

### Isolation and culturing of HSPCs

Hematopoietic stem and progenitor cells (HSPCs) were isolated from umbilical cord blood (Donor 1), obtained from the New York Blood Center, as previously described^8^. Mononuclear cells were separated by Ficoll-Paque (Cytiva, cat# 17144002) density gradient centrifugation and either cryopreserved in 7.5% bovine serum albumin (BSA, Life Technologies, cat# 501215315) and 10% DMSO (Hybri-Max, Sigma-Aldrich, cat# D2650) in PBS or processed immediately. For HSPC enrichment, cord blood mononuclear cells were incubated in Stemspan SFEM II (StemCell Technologies, cat# 09655) for 2 hours to remove adherent cells, after which nonadherent cells were sorted for CD133+ cells using a CD133 MicroBead kit (Miltenyi Biotech, cat# 130100830) protocol.

Additionally, anonymized presorted HSPCs were obtained from AllCells (cat# mLP RegH) as mobilized CD34+ cells (Donor 2). Cells were thawed in RPMI-1640 containing 10 mM HEPES (Invitrogen, cat# 15630080) and DNase I (Roche, cat# 04716728001).

HSPCs were cultured in STIF medium, Stemspan SFEM II supplemented with CC110 cytokine cocktail containing stem cell factor (SCF), Flt3 ligand (Flt3L), and thrombopoietin (TPO), each at 100 ng/mL (Stemcell Technologies, cat# 02697), and 100 ng/mL recombinant IGFBP2 (R&D Systems, cat# 674-B2-025). Cells were maintained at 37°C for 4 days before infection via spinoculation.

### Construction of dual-color HIV reporter constructs VT1 and VT1-IRES-cDNA

HIV-1 89.6-based dual-color latency reporter VT1 and VT1-IRES-MCS were constructed as previously described^34^. For VT1-IRES-cDNA reporter construct variants, PCR amplicons of BFP, GFI1, BCL2, EZR, TNFSF13B, ITGB2, or TRIM22 were cloned into VT1-IRES-MCS using AgeI and PmeI. To create VT1-IRES-ZFP36L2, ZFP36L2 was cloned into VT1-IRES-MCS via HpaI and PmeI. All gene cDNAs were MGC Human sequence-verified clones purchased from Horizon Discovery.

Oligonucleotide primers used for PCR amplification: AgeI-BFP-forward, AAAACCGGTATG AGCGAGCTGATTAAGGAGAACATG; PmeI-BFP-reverse, AAAGTTTAAACTCAATTAAGCTTGTGCCCCAGTTTG; AgeI-GFI1-forward, AAAACCGGTATGCCGCGCTCATT TCTCGTC; PmeI-GFI1-reverse, TTTGTTTAAACTCATTTGAGCCCATGCTGCGTC; HpaI-ZFP36L2-forward, AAAGTTAACATGTCGACCACACTTCTGTCCG; PmeI-ZFP36L2-reverse, TTTGTTTAAACTCAGTCGTCGGAGATGGAGAGG; AgeI-TNFSF13B-forward, AAAACCGGTATGGATGACTCCACAGAAAGGG; PmeI-TNFSF13B-reverse, AAAGTTTAAACTCACAGCAGTTTCAATGCACC; AgeI-ITGB2-forward, AAAACCGGTATG CTGGGCCTGCGC; PmeI-ITGB2-reverse, AAAGTTTAAACCTAACT CTCAGCAAACTTGGG GTTCATG; AgeI-EZR-forward, AAAACCGGTATGCCG AAACCAATCAATGTCC; PmeI-EZR-reverse, AAAGTTTAAACTTACAGGGCCTCG AACTCG; AgeI-BCL2-forward, AAAACCGGTATGGCGCACGCTGGG; PmeI-BCL2-reverse, AAAGTTTAAACTCACTTGTGGCCCAGATAGGC; AgeI-TRIM22-forward, AAAACCGGTATGGATTTC TCAGTAAAGGTAGACATAGAG; PmeI-TRIM22-reverse, AAAGTTTAAACTCAGGAGCTCGGTGGGC.

### Cloning of VT1-hU6-shRNA Reporter Constructs

The VT1-hU6-MCS vector was generated by synthesizing an hU6-MCS gBlock fragment and cloning it into VT1 using XbaI and PmeI restriction sites. Short hairpin RNAs (shRNAs) targeting GFI1, as well as scrambled controls, were designed using the Broad Institute Genetic Perturbation Platform. Annealed oligonucleotides encoding the shRNA hairpin (one forward and one reverse for each target) were cloned into the AgeI and ClaI sites of VT1-hU6-MCS to form VT1-hU6-shRNA. Oligonucleotide sequences used for shRNA constructs are listed below. The sequence of the synthesized hU6-MCS gBlocks DNA fragment (IDT) (XbaI-hU6-MCS-PmeI, 296 bp) is:

AAATCTAGAGAGGGCCTATTTCCCATGATTCCTTCATATTTGCATATACGATACAAGGCTGTTAGAGAGATAATTGGAATTAATTTGACTGTAAACACAAAGATATTAGTACAAAATACGTGACGTAGAAAGTAATAATTTCTTGGGTAGTTTGCAGTTTTAAAATTATGTTTTAAAATGGACTATCATATGCTTACCGTAACTTGAAAGTATTTCGATTTCTTGGCTTTATATATCTTGTGGAAAGGACGAAACACCGGTCCGCAGGTATGCACGCGTGTTAACGTTTAAACAA A Oligonucleotide sequences used for shRNA constructs: shRNA-74, ATTCCCTCCGTTTAGAGAATG; shRNA-978, ACTGTCCACACACCTGCTTAT; shRNA-1052, AGAAGTCAGACATGAAGAAAC; shRNA-1060, GACATGAAGAAACACACTTTC; shRNA-1247, AGACGCAGCATGGGCTCAAAT; shRNA-1834, AGGTGGACAAATGCTAATATT.

### Construction of NL4-3 BR3-1 dual-color reporter

pNL4-3 ΔGPE-GFP (McNamara et al., 2012^8^) was digested with XhoI (NEB cat# R0146) and NaeI (NEB cat# R0190), followed by dephosphorylation and gel purification. A synthetic 2939 bp gene fragment (Genscript GenTitan) containing 15 bp of overlap with the digested vector, a constitutive spleen focus-forming virus promoter (pSFFV) with Kozak sequence driving mCherry, and a 3’LTR bearing a mutated polyA signal and SV40 polyA signal was generated. Vector and insert were assembled using the Geneart Seamless Cloning kit (Fisher, cat# A14606), following the manufacturer’s protocol.

### Construction of pcDNA 3.1 expression constructs

pcDNA3.1-3xFlag-GFI1 was generated by cloning a GFI1 PCR amplicon (MGC Human sequence-verified cDNA, Horizon Discovery) into the BamHI and NotI sites of the pcDNA3.1-3xFlag vector using forward primer AAAGGATCCACCATGCCGCGCTCATTTCTC and reverse primer AAAGCGGCCGCTCATTTGAGCCCATGCTGC. A version without the Flag tag (pcDNA-GFI1) was created by deleting the N-terminal Flag sequence using the Q5 Site-Directed Mutagenesis kit (New England Biolabs, cat# E0554) and exclusion PCR (forward primer ATG CCGCGCTCATTTCTC; reverse primer GCTAGCCAGCTTGGGTCT). The GFI1 N382S mutant was generated via site-directed mutagenesis (Q5 kit) using forward primer CAGAGCTCCAGCCTCATCACC and reverse primer GCTGAATGCCTTGCCGCA, and the P2A mutation was introduced similarly, with forward primer GGCTAGCATGGCTCGCTCATTTC and reverse primer AGCTTGGGTCTCCCTATAG. All mutagenesis primers were designed using the NEBaseChanger tool (https://nebasechanger.neb.com/) and reactions followed the Q5 Site-Directed Mutagenesis kit manufacturer’s instructions.

The pcDNA-3.1 3X Flag-ZFP961 construct was kindly provided by Sherry Ralls (NIH/NICHD/PGD/SMER, Bethesda, MD). The pcDNA-3.1 3X Flag-ZFP809(1–353) plasmid was a gift from Stephen Goff (Addgene plasmid # 67788). The ZNF627 open reading frame (GenScript, OHu0653OD0; NM_145295.4) was supplied pre-cloned into the pcDNA3.1(+)/C-(K)-DYK vector (GenScript), resulting in a construct with a C-terminal FLAG tag. Separately, the ZFP627 coding sequence was excised from the GenScript plasmid using KpnI and XhoI digestion and ligated into the pcDNA3.1 3X-FLAG vector, resulting in an N-terminal FLAG-tagged ZFP627 construct.

### Virus production

HEK293T cells were seeded at 2.5 × 10⁶ per 10 cm dish and allowed to adhere overnight. Cells were transfected the next day with 12 µg total plasmid DNA, comprising equal amounts of pCMV-HIV (S.-J.-K. Yee, City of Hope National Medical Center), pHCMV-G (VSV-G envelope; Nancy Hopkins, MIT), and the lentiviral expression plasmid, using polyethylenimine (PEI; Polysciences, cat# 23966-2) at a 4:1 PEI:DNA ratio. Plasmids were diluted in 150 mM NaCl, combined with PEI, vortexed, and incubated at room temperature for 15 minutes before addition to the cells. 8 hrs post-transfection, the medium was replaced with fresh D10 as mentioned above. Viral supernatant was collected 48 hours later, centrifuged to remove debris, aliquoted, and stored at –80°C.

### Viral transductions

Virus was diluted in cell culture medium to achieve the desired infection rate, determined by serial dilution. CEM-A2 cells were resuspended and plated at 250,000 cells per well in 48-well plates. To enhance transduction efficiency, spinoculation was performed at 1,050 × g for 2 hours at 25°C. Following spinoculation, the medium was aspirated and replaced with fresh R10 medium. For VT1-IRES cDNA overexpression experiments, cells were incubated for 5 days post-transduction. For VT1-hU6-shRNA knockdown experiments, transduced cells were harvested 7 days post-transduction for flow cytometry or sorted for western blot analysis.

For HSPCs, cells were transduced with virus at a density of 500,000 cells/mL in 24-well plates with virus. Polybrene (Millipore Sigma, cat# H9268-5G) at 4 µg/mL was added to the virus-cell mixture. The cells were then spinoculated under the same conditions as above. After centrifugation, the medium was aspirated and replaced with STIF medium. HSPC cultures were incubated at either 37°C or 30°C with 5% CO₂ for 3 days prior to sorting for single-cell RNA sequencing.

### Western blotting

For western blot analysis, cells were lysed in Blue Loading Buffer (cat# 7722, Cell Signaling Technology), sonicated with a Misonix sonicator (Qsonica, LLC. Newtown, CT), incubated in a heating block for 10 min at 95°C before loading, and analyzed by SDS-PAGE immunoblot. Samples were loaded on a 4-15% Criterion TGX precast midi protein gel (Bio-Rad, cat# 5671083) and ran at 130 V for 120 min. Protein was transferred to a PVDF membrane (Millipore, cat# IPVH00005) at 350 mAmp for 90 minutes. The membrane was blocked with 5% milk (Fisher, cat# NC9121673) in TBST (0.05% Tween-20 (Fisher, cat# BP337-500), 150 mM sodium chloride (NaCl, Fisher, cat# BP358-212), 10 mM Tris(hydroxymethyl)aminomethane (Tris, Invitrogen, cat# 15504020), pH 8) for 1 hr at room temperature with rocking. The blot was then probed with the antibodies indicated below and imaged with a ChemiDoc Imaging System (BioRad, cat# 12003153).

Antibodies: mouse anti-vinculin (1:1000, Cat# V9131, Millipore Sigma), mouse anti-GFI1 (1:1000; or 1:250 for knockdown experiments, Cat# WH0002672M1, Sigma), chicken anti-GFP (1:2000, Cat# 13970, Abcam). Secondary antibodies included HRP-conjugated goat anti-mouse IgG (1:10,000, Cat# 31430, Invitrogen) and HRP-conjugated goat anti-chicken IgY (1:10,000, Cat# A16054, Invitrogen).

### EMSA

#### Nuclear extraction

HEK293T cells were plated at 7.5 ×10⁶ per 15 cm dish and incubated for 24 hours. Cells were transfected with 30 µg DNA in 150 mM NaCl using PEI 4 μL of PEI (1 mg/mL) per 1 μg of plasmid DNA. After 48 hours, cells were collected by pelleting, washed, and resuspended in PBS. Cells were pelleted again and lysed in Dignam buffer A (10 mM HEPES, pH 7.9, 1.5 mM MgCl₂, 10 mM KCl, 0.5 mM DTT, Halt protease inhibitor) (ThermoFisher, cat# 87786 by performing 15 strokes with a 2 mL Dounce homogenizer (cat# 501945204).

Nuclei were pelleted by centrifugation at 4300 × g for 5 minutes, and the supernatant removed. The nuclei were resuspended in Dignam buffer C (20 mM HEPES, pH 7.9, 1.5 mM MgCl₂, 25% v/v glycerol, 0.2 mM EDTA, 0.5 mM DTT, Halt protease inhibitor) and lysed by 10 strokes with the 2 mL Dounce homogenizer. Nuclear extracts were clarified by centrifugation at 21,130 × g for 15 minutes, and the supernatants were aliquoted, snap-frozen, and stored at –80°C.

#### Oligo annealing

The probes, as shown in Fig. 4A, were synthesized as single-stranded oligonucleotides, each labeled at the 5’ end with IRDye 700 (IDT). To prepare duplex probes, 5μL of 20pmol/μL forward IRDye 700 oligo and reverse IRDye 700 oligo were mixed and annealed at 100°C for 4 minutes, followed by 78 cycles, each with a 1°C decrease and 1 minute per cycle. The annealed oligonucleotide probes were diluted to a 100 fmol/μL working stock and stored at –20°C.

250fmol of probes were incubated with nuclear extract in modified Thornell binding buffer (25 mM HEPES, pH 7.9, 1 mM EDTA, 10% v/v glycerol, 5 mM DTT, 5 mM NaCl, 5 mM KCl, 3 mM MgCl₂, 0.1 mM ZnCl₂) for 30 minutes at 30°C in a total volume of 20 μL. Poly dI-dC (1 μg per 20 μL reaction; ThermoFisher, cat# PI20148E) was used as a nonspecific competitor and pre-incubated with nuclear lysates for 20 minutes before addition of EMSA probes. For competition experiments, a 500-fold molar excess of cold (unlabeled) probe was added. Samples were loaded with orange loading dye (Li-Cor, cat# 92710100) onto 5% polyacrylamide TBE gels (Bio-Rad, cat#4565014) and run in 0.5× TBE buffer for 45 minutes at 100 V protected from light. Gels were imaged using an Amersham Typhoon 5 imager (Cytiva, cat#29187194).

### HEK293T co-transfections

HEK293T cells were plated at 125,000 cells per well in 12-well plates and allowed to adhere overnight. The next day, the HIV reporter construct of interest and pcDNA3.1 expression construct containing the gene of interest were diluted in 150 mM NaCl. Reporter constructs VT1, VT2, and BR3-1 were transfected to achieve a transfection rate of approximately 15–30%. The pcDNA3.1 expression construct was used at 100 fmol per well, except where otherwise indicated in figure panels. To ensure equal total DNA per well, pUC19 carrier plasmid was added to adjust DNA quantities as needed.

PEI was added at a 4:1 mass ratio PEI:DNA, and the mixture was gently vortexed for 15 seconds. The transfection solution was incubated at room temperature for 15 minutes before being added to each well. Cells were incubated for 24 hours post-transfection prior to flow cytometry analysis or immunoblotting.

### NF-κB stimulation and inhibition treatments

HEK293T cells were co-transfected with the VT2 LTR reporter plasmid and either pcDNA-EV or GFI1. For NF-κB activation, cells were treated with TNF-α (R&D Systems, cat# 210-TA-005). To inhibit NF-κB activation, cells were co-treated with an IKK inhibitor (IKK2 inhibitor VI, Cayman Chemical, cat# 17276). Treatment concentrations and timing are provided in the figure panels.

### Combination LRA treatment

CEM-A2 cells transduced with VT1-IRES-GFI1 or BFP control virus were cultured for 4 days, then treated with latency-reversing agents (LRAs) 24 hours prior to harvest and cytometric analysis. The following LRAs were used: 1 µM entinostat (Selleckchem, cat# S1053), 5 nM bryostatin-1 (Sigma-Aldrich, cat# B7431), 5 µM GSK-LSD1 (Selleckchem, cat# S7574), and 1 µM A366 (Selleckchem, cat# S7572). Treatments were administered individually or in combinations as indicated in Fig. 8.

For HEK293T cell compound treatments, cells were co-transfected with 100 fmol pcDNA-GFI1 or EV and VT2 reporter plasmid. 24 hours post-transfection, cells were treated with 1 µM entinostat, 5 µM GSK-LSD1, and 1 µM A366, and collected 24 hours later for flow cytometry analysis.

### Flow cytometry and fluorescence-activated cell sorting (FACS)

Protein expression in transduced CEM-A2 cells and transiently transfected HEK293T cells was assessed using a Cytek Aurora flow cytometer. In all experiments, cells were initially gated by forward scatter versus side scatter, followed by doublet exclusion. Subsequent gating was performed first on constitutive reporter fluorescence and then on LTR reporter fluorescence. Cytometric data were analyzed with FlowJo v10 software (BD Life Sciences).

For HSPCs, actively infected (GFP⁺/mCherry⁺) and latently infected (GFP⁺/mCherry⁻) cells were individually sorted using a Sony SH800 cell sorter. Active and latent cell populations were then combined at a 1:1 ratio for single-cell RNA sequencing.

### Integration of scRNA-seq datasets

LIGER^43,44^ was used to integrate the four HIV-exposed scRNA-seq samples to remove batch effects. The four samples are Donor 1 and Donor 2 at 30℃ or 37℃. As an initial quality control step, cells were filtered with criteria of number of UMIs > 1,000, number of genes > 100, mitochondrial percentage < 10, and ribosomal RNA percentage > 10. HIV, mCherry, and EGFP expressions were not included during integration. The integration was run with k=20 and quantileNorm. Clusters were found with resolution of 0.2 using LIGER runCluster function, and UMAP embedding was generated with cosine distance metric, 30 neighbors, and 0.3 minimum distance using LIGER runUMAP function. We labeled cell types based on canonical blood marker genes, including *SPINK2*, *MPO*, *LYZ*, *TCF4*, *HBD*, *HDC*, *RHEX*, *PF4*, and *EBF1*.

### Latency classification

We extracted HIV expression values (labeled as “HIV” in the feature matrix) for each sample and applied a log(x+1) transformation. To distinguish between active (high expression) and latent (low expression) populations, we manually defined a cutoff threshold located near the local minimum of each sample’s density distribution. The cutoff values were capped between 4 and 6 to ensure the split reflected the nearly balanced representation established during sample preparation, where latently infected and actively infected populations were combined in a 1:1 ratio post FACS sorting.

### Differential gene expression analysis

Following the LIGER integration and cell latency annotation of the four samples, differentially expressed markers between active and latent populations were found using LIGER runMarkerDEG function with Wilcoxon rank-sum test. The volcano plot depicts genes with differential expression patterns in active (left) or latent (right) cells. We selected logFC threshold of 0.5 and p-adjusted value threshold of 0.05 as cutoffs. Important latency-associated genes are annotated and plotted as red dots.

### Scatterplot of HIV and human gene expression

To mitigate the sparse expression issue of single-cell sequencing data, the plotted HIV expression values and human gene expressions were first smoothed by cell neighbors in UMAP embedded space, specifically, the connectivity matrix stored in the HSPC AnnData object during Scanpy preprocessing steps was used to convolve the expression arrays, following a similar approach to scVelo’s pp.moments function. The scatter plots were generated for each of the four scRNA-seq samples.

### Statistical analysis

Statistical analyses for all experiments, except for single-cell data, were performed using GraphPad Prism v10 software (Boston, MA), as described in the figure legends. Analyses involving single-cell data were conducted separately, with details provided above.

## ACKNOWLEDGMENTS

This work was supported by National Institutes of Health grants R01AI149669 to K.L.C. and J.D.W. We thank the University of Michigan Flow Cytometry Core for providing access to instrumentation. Library preparation and single-cell RNA sequencing services were provided by the Advanced Genomics Core at the University of Michigan. We are also grateful to all members of the Collins lab for their helpful discussions. Cartoons in Figs. 1A, 3B, 3G, 4B, 5A, 7A, B, and D, 8A and D, and Fig. 9 were created with BioRender.com (2025).

## AUTHOR CONTRIBUTIONS

Conceptualization, J.A.O, C.C, W.M.D, M.C.V, M.M.P and J.D.W, K.L.C.; Methodology, J.A.O, C.L, C.C, W.M.D, M.C.V, M.M.P, V.T, J.D.W., and K.L.C.; Investigation, J.A.O,C.L,C.C, W.M.D.,M.M.P; Analysis, J.A.O, C.L, C.C, W.M.D., M.M.P. Resources J.A.O, C.L, C.C, W.M.D, V.T, B.R, J.D.W, K.L.C.; Writing – Original Draft, J.A.O.; Writing – Review & Editing, J.A.O., and K.L.C.; Supervision, J.D.W. and K.L.C., Funding Acquisition, J.D.W., and K.L.C.

## REFERENCES

1. Deeks, S. G. et al. Research priorities for an HIV cure: International AIDS Society Global Scientific Strategy 2021. Nat. Med. 27, 2085–2098 (2021).

2. Chun, T.-W. et al. Early establishment of a pool of latently infected, resting CD4+ T cells during primary HIV-1 infection. Proc. Natl. Acad. Sci. 95, 8869–8873 (1998).

3. Finzi, D. et al. Identification of a reservoir for HIV-1 in patients on highly active antiretroviral therapy. Science 278, 1295–1300 (1997).

4. Pitman, M. C., Lau, J. S. Y., McMahon, J. H. & Lewin, S. R. Barriers and strategies to achieve a cure for HIV. Lancet HIV 5, e317–e328 (2018).

5. Williams, J. P. et al. HIV-1 DNA predicts disease progression and post-treatment virological control. eLife 3, e03821.

6. Ho, Y.-C. et al. Replication-Competent Noninduced Proviruses in the Latent Reservoir Increase Barrier to HIV-1 Cure. Cell 155, 540–551 (2013).

7. Carter, C. C. et al. HIV-1 infects multipotent progenitor cells causing cell death and establishing latent cellular reservoirs. Nat. Med. 16, 446–451 (2010).

8. McNamara, L. A., Ganesh, J. A. & Collins, K. L. Latent HIV-1 infection occurs in multiple subsets of hematopoietic progenitor cells and is reversed by NF-κB activation. J. Virol. 86, 9337–9350 (2012).

9. Zaikos, T. D. et al. Hematopoietic Stem and Progenitor Cells Are a Distinct HIV Reservoir that Contributes to Persistent Viremia in Suppressed Patients. Cell Rep. 25, 3759–3773.e9 (2018).

10. Orkin, S. H. & Zon, L. I. Hematopoiesis: an evolving paradigm for stem cell biology. Cell 132, 631–644 (2008).

11. Durdik, M. et al. Hematopoietic stem/progenitor cells are less prone to undergo apoptosis than lymphocytes despite similar DNA damage response. Oncotarget 8, 48846–48853 (2017).

12. Painter, M. M., Zaikos, T. D. & Collins, K. L. Quiescence Promotes Latent HIV Infection and Resistance to Reactivation from Latency with Histone Deacetylase Inhibitors. J. Virol. 91, e01080–17 (2017).

13. Zaikos, T. D., Painter, M. M., Sebastian Kettinger, N. T., Terry, V. H. & Collins, K. L. Class 1-Selective Histone Deacetylase (HDAC) Inhibitors Enhance HIV Latency Reversal while Preserving the Activity of HDAC Isoforms Necessary for Maximal HIV Gene Expression. J. Virol. 92, e02110–17 (2018).

14. Carter, C. C. et al. HIV-1 utilizes the CXCR4 chemokine receptor to infect multipotent hematopoietic stem and progenitor cells. Cell Host Microbe 9, 223–234 (2011).

15. Sebastian, N. T. et al. CD4 is expressed on a heterogeneous subset of hematopoietic progenitors, which persistently harbor CXCR4 and CCR5-tropic HIV proviral genomes in vivo. PLoS Pathog. 13, e1006509 (2017).

16. Han, Y. et al. Resting CD4+ T cells from human immunodeficiency virus type 1 (HIV-1)-infected individuals carry integrated HIV-1 genomes within actively transcribed host genes. J. Virol. 78, 6122–6133 (2004).

17. Schröder, A. R. W. et al. HIV-1 Integration in the Human Genome Favors Active Genes and Local Hotspots. Cell 110, 521–529 (2002).

18. Friedman, J. et al. Epigenetic silencing of HIV-1 by the histone H3 lysine 27 methyltransferase enhancer of Zeste 2. J. Virol. 85, 9078–9089 (2011).

19. Imai, K., Togami, H. & Okamoto, T. Involvement of histone H3 lysine 9 (H3K9) methyltransferase G9a in the maintenance of HIV-1 latency and its reactivation by BIX01294. J. Biol. Chem. 285, 16538–16545 (2010).

20. Pearson, R. et al. Epigenetic silencing of human immunodeficiency virus (HIV) transcription by formation of restrictive chromatin structures at the viral long terminal repeat drives the progressive entry of HIV into latency. J. Virol. 82, 12291–12303 (2008).

21. Chéné, I. du et al. Suv39H1 and HP1γ are responsible for chromatin-mediated HIV-1 transcriptional silencing and post-integration latency. EMBO J. 26, 424–435 (2007).

22. Blazkova, J. et al. CpG methylation controls reactivation of HIV from latency. PLoS Pathog. 5, e1000554 (2009).

23. Kauder, S. E., Bosque, A., Lindqvist, A., Planelles, V. & Verdin, E. Epigenetic regulation of HIV-1 latency by cytosine methylation. PLoS Pathog. 5, e1000495 (2009).

24. Wolf, G. et al. The KRAB zinc finger protein ZFP809 is required to initiate epigenetic silencing of endogenous retroviruses. Genes Dev. 29, 538–554 (2015).

25. Wolf, G. et al. KRAB-zinc finger protein gene expansion in response to active retrotransposons in the murine lineage. eLife 9, e56337 (2020).

26. Yang, B., et al. Species-specific KRAB-ZFPs function as repressors of retroviruses by targeting PBS regions. Proc. Natl. Acad. Sci. 119, e2119415119 (2022).

27. Wolf, D. & Goff, S. P. TRIM28 Mediates Primer Binding Site-Targeted Silencing of Murine Leukemia Virus in Embryonic Cells. Cell 131, 46–57 (2007).

28. Perez, M. et al. Bryostatin-1 Synergizes with Histone Deacetylase Inhibitors to Reactivate HIV-1 from Latency. Curr. HIV Res. 8, 418–429.

29. Nabel, G. & Baltimore, D. An inducible transcription factor activates expression of human immunodeficiency virus in T cells. Nature 326, 711–713 (1987).

30. Williams, S. A., Kwon, H., Chen, L.-F. & Greene, W. C. Sustained induction of NF-kappa B is required for efficient expression of latent human immunodeficiency virus type 1. J. Virol. 81, 6043–6056 (2007).

31. Barboric, M. et al. Tat competes with HEXIM1 to increase the active pool of P-TEFb for HIV-1 transcription. Nucleic Acids Res. 35, 2003–2012 (2007).

32. Parada, C. A. & Roeder, R. G. Enhanced processivity of RNA polymerase II triggered by Tat-induced phosphorylation of its carboxy-terminal domain. Nature 384, 375–378 (1996).

33. Balboni, P. G. et al. Inhibition of human immunodeficiency virus reactivation from latency by a tat transdominant negative mutant. J. Med. Virol. 41, 289–295 (1993).

34. Gomez-Rivera, F. et al. Variation in HIV-1 Tat activity is a key determinant in the establishment of latent infection. JCI Insight e184711 (2024) doi:10.1172/jci.insight.184711.

35. Contreras, X. et al. Suberoylanilide Hydroxamic Acid Reactivates HIV from Latently Infected Cells. J. Biol. Chem. 284, 6782–6789 (2009).

36. Wei, D. G. et al. Histone deacetylase inhibitor romidepsin induces HIV expression in CD4 T cells from patients on suppressive antiretroviral therapy at concentrations achieved by clinical dosing. PLoS Pathog. 10, e1004071 (2014).

37. Contreras, X., Barboric, M., Lenasi, T. & Peterlin, B. M. HMBA Releases P-TEFb from HEXIM1 and 7SK snRNA via PI3K/Akt and Activates HIV Transcription. PLOS Pathog. 3, e146 (2007).

38. Vlach, J. & Pitha, P. M. Hexamethylene bisacetamide activates the human immunodeficiency virus type 1 provirus by an NF-κB-independent mechanism. J. Gen. Virol. 74, 2401–2408 (1993).

39. Verdin, E., Paras, P. & Van Lint, C. Chromatin disruption in the promoter of human immunodeficiency virus type 1 during transcriptional activation. EMBO J. 12, 3249–3259 (1993).

40. Rafati, H. et al. Repressive LTR nucleosome positioning by the BAF complex is required for HIV latency. PLoS Biol. 9, e1001206 (2011).

41. Nguyen, K., Das, B., Dobrowolski, C. & Karn, J. Multiple Histone Lysine Methyltransferases Are Required for the Establishment and Maintenance of HIV-1 Latency. mBio 8, e00133–17 (2017).

42. Tyagi, M., Pearson, R. J. & Karn, J. Establishment of HIV latency in primary CD4+ cells is due to epigenetic transcriptional silencing and P-TEFb restriction. J. Virol. 84, 6425–6437 (2010).

43. Welch, J. D. et al. Single-Cell Multi-omic Integration Compares and Contrasts Features of Brain Cell Identity. Cell 177, 1873–1887.e17 (2019).

44. Liu, J. et al. Jointly defining cell types from multiple single-cell datasets using LIGER. Nat. Protoc. 15, 3632–3662 (2020).

45. Sharif-Askari, E. et al. Zinc finger protein Gfi1 controls the endotoxin-mediated Toll-like receptor inflammatory response by antagonizing NF-kappaB p65. Mol. Cell. Biol. 30, 3929–3942 (2010).

46. Guo, G. et al. Gfi1 and Zc3h12c orchestrate a negative feedback loop that inhibits NF-kB activation during inflammation in macrophages. Mol. Immunol. 128, 219–226 (2020).

47. Liu, P., Yang, P., Zhang, Z., Liu, M. & Hu, S. Ezrin/NF-κB Pathway Regulates EGF-induced Epithelial-Mesenchymal Transition (EMT), Metastasis, and Progression of Osteosarcoma. Med. Sci. Monit. Int. Med. J. Exp. Clin. Res. 24, 2098–2108 (2018).

48. Makita, S., Takatori, H. & Nakajima, H. Post-Transcriptional Regulation of Immune Responses and Inflammatory Diseases by RNA-Binding ZFP36 Family Proteins. Front. Immunol. 12, (2021).

49. Yee, N. K. & Hamerman, J. A. β(2) integrins inhibit TLR responses by regulating NF-κB pathway and p38 MAPK activation. Eur. J. Immunol. 43, 779–792 (2013).

50. Grimm, S., Bauer, M. K., Baeuerle, P. A. & Schulze-Osthoff, K. Bcl-2 down-regulates the activity of transcription factor NF-kappaB induced upon apoptosis. J. Cell Biol. 134, 13–23 (1996).

51. Meng, L. et al. S100A9 Derived From Myeloma Associated Myeloid Cells Promotes TNFSF13B/TNFRSF13B-Dependent Proliferation and Survival of Myeloma Cells. Front. Oncol. 11, 691705 (2021).

52. Ji, J. et al. TRIM22 activates NF-κB signaling in glioblastoma by accelerating the degradation of IκBα. Cell Death Differ. 28, 367–381 (2021).

53. Roeth, J. F., Williams, M., Kasper, M. R., Filzen, T. M. & Collins, K. L. HIV-1 Nef disrupts MHC-I trafficking by recruiting AP-1 to the MHC-I cytoplasmic tail. J. Cell Biol. 167, 903–913 (2004).

54. Zweidler-Mckay, P. A., Grimes, H. L., Flubacher, M. M. & Tsichlis, P. N. Gfi-1 encodes a nuclear zinc finger protein that binds DNA and functions as a transcriptional repressor. Mol. Cell. Biol. 16, 4024–4034 (1996).

55. Duan, Z. & Horwitz, M. Targets of the transcriptional repressor oncoprotein Gfi-1. Proc. Natl. Acad. Sci. 100, 5932–5937 (2003).

56. Grimes, H. L., Gilks, C. B., Chan, T. O., Porter, S. & Tsichlis, P. N. The Gfi-1 protooncoprotein represses Bax expression and inhibits T-cell death. Proc. Natl. Acad. Sci. 93, 14569–14573 (1996).

57. Gilboa, E., Mitra, S. W., Goff, S. & Baltimore, D. A detailed model of reverse transcription and tests of crucial aspects. Cell 18, 93–100 (1979).

58. Harada, F., Peters, G. G. & Dahlberg, J. E. The primer tRNA for Moloney murine leukemia virus DNA synthesis. Nucleotide sequence and aminoacylation of tRNAPro. J. Biol. Chem. 254, 10979–10985 (1979).

59. Li, X. et al. Effects of alterations of primer-binding site sequences on human immunodeficiency virus type 1 replication. J. Virol. 68, 6198–6206 (1994).

60. Mak, J. & Kleiman, L. Primer tRNAs for reverse transcription. J. Virol. 71, 8087–8095 (1997).

61. Miller, S. B., Yildiz, F. Z., Lo, J. A., Wang, B. & D’Souza, V. M. A structure-based mechanism for tRNA and retroviral RNA remodelling during primer annealing. Nature 515, 591–595 (2014).

62. Wolf, D. & Goff, S. P. Embryonic stem cells use ZFP809 to silence retroviral DNAs. Nature 458, 1201–1204 (2009).

63. Collman, R. et al. An infectious molecular clone of an unusual macrophage-tropic and highly cytopathic strain of human immunodeficiency virus type 1. J. Virol. 66, 7517–7521 (1992).

64. Adachi, A. et al. Production of acquired immunodeficiency syndrome-associated retrovirus in human and nonhuman cells transfected with an infectious molecular clone. J. Virol. 59, 284–291 (1986).

65. Thambyrajah, R. et al. GFI1 proteins orchestrate the emergence of haematopoietic stem cells through recruitment of LSD1. Nat. Cell Biol. 18, 21–32 (2016).

66. Saleque, S., Kim, J., Rooke, H. M. & Orkin, S. H. Epigenetic Regulation of Hematopoietic Differentiation by Gfi-1 and Gfi-1b Is Mediated by the Cofactors CoREST and LSD1. Mol. Cell 27, 562–572 (2007).

67. Grimes, H. L., Chan, T. O., Zweidler-McKay, P. A., Tong, B. & Tsichlis, P. N. The Gfi-1 proto-oncoprotein contains a novel transcriptional repressor domain, SNAG, and inhibits G1 arrest induced by interleukin-2 withdrawal. Mol. Cell. Biol. 16, 6263–6272 (1996).

68. Möröy, T. & Khandanpour, C. Role of GFI1 in Epigenetic Regulation of MDS and AML Pathogenesis: Mechanisms and Therapeutic Implications. Front. Oncol. 9, (2019).

69. Duan, Z., Zarebski, A., Montoya-Durango, D., Grimes, H. L. & Horwitz, M. Gfi1 Coordinates Epigenetic Repression of p21Cip/WAF1 by Recruitment of Histone Lysine Methyltransferase G9a and Histone Deacetylase 1. Mol. Cell. Biol. 25, 10338–10351 (2005).

70. McGhee, L. et al. Gfi-1 attaches to the nuclear matrix, associates with ETO (MTG8) and histone deacetylase proteins, and represses transcription using a TSA-sensitive mechanism. J. Cell. Biochem. 89, 1005–1018 (2003).

71. Gilks, C. B., Bear, S. E., Grimes, H. L. & Tsichlis, P. N. Progression of interleukin-2 (IL-2)-dependent rat T cell lymphoma lines to IL-2-independent growth following activation of a gene (Gfi-1) encoding a novel zinc finger protein. Mol. Cell. Biol. 13, 1759–1768 (1993).

72. Lee, S. et al. Solution structure of Gfi-1 zinc domain bound to consensus DNA. J. Mol. Biol. 397, 1055–1066 (2010).

73. Person, R. E. et al. Mutations in proto-oncogene GFI1 cause human neutropenia and target ELA2. Nat. Genet. 34, 308–312 (2003).

74. Zarebski, A. P., Baktula, A. M., Basu, S., Trent, J. O. & Grimes, H. L. Gfi1-N382S Mutants from Human Severe Congenital Neutropenia Patients Function through a Transcriptional Dominant Negative Mechanism. Blood 108, 1185 (2006).

75. Zarebski, A. et al. Mutations in Growth Factor Independent-1 Associated with Human Neutropenia Block Murine Granulopoiesis through Colony Stimulating Factor-1. Immunity 28, 370–380 (2008).

76. Sedore, S. C. et al. Manipulation of P-TEFb control machinery by HIV: recruitment of P-TEFb from the large form by Tat and binding of HEXIM1 to TAR. Nucleic Acids Res. 35, 4347–4358 (2007).

77. Zhu, Y. et al. Transcription elongation factor P-TEFb is required for HIV-1 Tat transactivation in vitro. Genes Dev. 11, 2622–2632 (1997).

78. Wei, P., Garber, M. E., Fang, S.-M., Fischer, W. H. & Jones, K. A. A Novel CDK9-Associated C-Type Cyclin Interacts Directly with HIV-1 Tat and Mediates Its High-Affinity, Loop-Specific Binding to TAR RNA. Cell 92, 451–462 (1998).

79. Hottiger, M. O. & Nabel, G. J. Interaction of Human Immunodeficiency Virus Type 1 Tat with the Transcriptional Coactivators p300 and CREB Binding Protein. J. Virol. 72, 8252–8256 (1998).

80. Marzio, G., Tyagi, M., Gutierrez, M. I. & Giacca, M. HIV-1 Tat transactivator recruits p300 and CREB-binding protein histone acetyltransferases to the viral promoter. Proc. Natl. Acad. Sci. 95, 13519–13524 (1998).

81. Benkirane, M. et al. Activation of Integrated Provirus Requires Histone Acetyltransferase: p300 AND P/CAF ARE COACTIVATORS FOR HIV-1 Tat *. J. Biol. Chem. 273, 24898–24905 (1998).

82. Petrusca, D. N. et al. Growth factor independence 1 expression in myeloma cells enhances their growth, survival, and osteoclastogenesis. J. Hematol. Oncol.J Hematol Oncol 11, 123 (2018).

83. Gerritsen, M. E. et al. CREB-binding protein/p300 are transcriptional coactivators of p65. Proc. Natl. Acad. Sci. 94, 2927–2932 (1997).

84. Perkins, N. D. et al. Regulation of NF-κB by Cyclin-Dependent Kinases Associated with the p300 Coactivator. Science 275, 523–527 (1997).

85. Mbonye, U. & Karn, J. The cell biology of HIV-1 latency and rebound. Retrovirology 21, 6 (2024).

86. Baxter, A. et al. Hit-to-lead studies: the discovery of potent, orally active, thiophenecarboxamide IKK-2 inhibitors. Bioorg. Med. Chem. Lett. 14, 2817–2822 (2004).

87. Li, Z. W. et al. The IKKbeta subunit of IkappaB kinase (IKK) is essential for nuclear factor kappaB activation and prevention of apoptosis. J. Exp. Med. 189, 1839–1845 (1999).

88. Guo, Q. et al. NF-κB in biology and targeted therapy: new insights and translational implications. Signal Transduct. Target. Ther. 9, 53 (2024).

89. Velinder, M. et al. GFI1 functions in transcriptional control and cell fate determination require SNAG domain methylation to recruit LSD1. Biochem. J. 473, 3355–3369 (2016).

90. Lee, C. et al. Lsd1 as a therapeutic target in Gfi1-activated medulloblastoma. Nat. Commun. 10, 332 (2019).

91. Park, S.-Y. & Kim, J.-S. A short guide to histone deacetylases including recent progress on class II enzymes. Exp. Mol. Med. 52, 204–212 (2020).

92. Khan, N. et al. Determination of the class and isoform selectivity of small-molecule histone deacetylase inhibitors. Biochem. J. 409, 581–589 (2007).

93. Hu, E. et al. Identification of Novel Isoform-Selective Inhibitors within Class I Histone Deacetylases. J. Pharmacol. Exp. Ther. 307, 720–728 (2003).

94. Barth, J. et al. LSD1 inhibition by tranylcypromine derivatives interferes with GFI1-mediated repression of PU.1 target genes and induces differentiation in AML. Leukemia 33, 1411–1426 (2019).

95. Shinkai, Y. & Tachibana, M. H3K9 methyltransferase G9a and the related molecule GLP. Genes Dev. 25, 781–788 (2011).

96. Tachibana, M., Sugimoto, K., Fukushima, T. & Shinkai, Y. Set domain-containing protein, G9a, is a novel lysine-preferring mammalian histone methyltransferase with hyperactivity and specific selectivity to lysines 9 and 27 of histone H3. J. Biol. Chem. 276, 25309–25317 (2001).

97. Mehla, R. et al. Bryostatin modulates latent HIV-1 infection via PKC and AMPK signaling but inhibits acute infection in a receptor independent manner. PloS One 5, e11160 (2010).

98. Raghuvanshi, R. & Bharate, S. B. Preclinical and Clinical Studies on Bryostatins, A Class of Marine-Derived Protein Kinase C Modulators: A Mini-Review. Curr. Top. Med. Chem. 20, 1124–1135 (2020).

99. McNamara, L. A. et al. CD133+ hematopoietic progenitor cells harbor HIV genomes in a subset of optimally treated people with long-term viral suppression. J. Infect. Dis. 207, 1807–1816 (2013).

100. van der Meer, L. T., Jansen, J. H. & van der Reijden, B. A. Gfi1 and Gfi1b: key regulators of hematopoiesis. Leukemia 24, 1834–1843 (2010).

101. Wang, C. & Goff, S. P. Differential control of retrovirus silencing in embryonic cells by proteasomal regulation of the ZFP809 retroviral repressor. Proc. Natl. Acad. Sci. 114, (2017).

102. Trono, D. et al. HIV Persistence and the Prospect of Long-Term Drug-Free Remissions for HIV-Infected Individuals. Science 329, 174–180 (2010).

103. Benkirane, M. et al. Activation of integrated provirus requires histone acetyltransferase. p300 and P/CAF are coactivators for HIV-1 Tat. J. Biol. Chem. 273, 24898–24905 (1998).

104. DeChristopher, B. A. et al. Designed, synthetically accessible bryostatin analogues potently induce activation of latent HIV reservoirs in vitro. Nat. Chem. 4, 705–710 (2012).

105. Kinter, A. L., Poli, G., Maury, W., Folks, T. M. & Fauci, A. S. Direct and cytokine-mediated activation of protein kinase C induces human immunodeficiency virus expression in chronically infected promonocytic cells. J. Virol. 64, 4306–4312 (1990).

106. Maiques-Diaz, A., Lynch, J. T., Spencer, G. J. & Somervaille, T. C. P. LSD1 inhibitors disrupt the GFI1 transcription repressor complex. Mol. Cell. Oncol. 5, e1481813 (2018).

107. Maiques-Diaz, A. et al. Enhancer Activation by Pharmacologic Displacement of LSD1 from GFI1 Induces Differentiation in Acute Myeloid Leukemia. Cell Rep. 22, 3641–3659 (2018).

108. Laird, G. M. et al. Ex vivo analysis identifies effective HIV-1 latency–reversing drug combinations. J. Clin. Invest. 125, 1901–1912 (2015).

109. Battistini, A. & Sgarbanti, M. HIV-1 latency: an update of molecular mechanisms and therapeutic strategies. Viruses 6, 1715–1758 (2014).

110. Fraszczak, J. & Möröy, T. The transcription factors GFI1 and GFI1B as modulators of the innate and acquired immune response. Adv. Immunol. 149, 35–94 (2021).

