## Supplementary figures for "Mechanisms of HIV Latency in Hematopoietic Progenitors: GFI1 as a Key Regulator"

### SUPPLEMENTARY INFORMATION

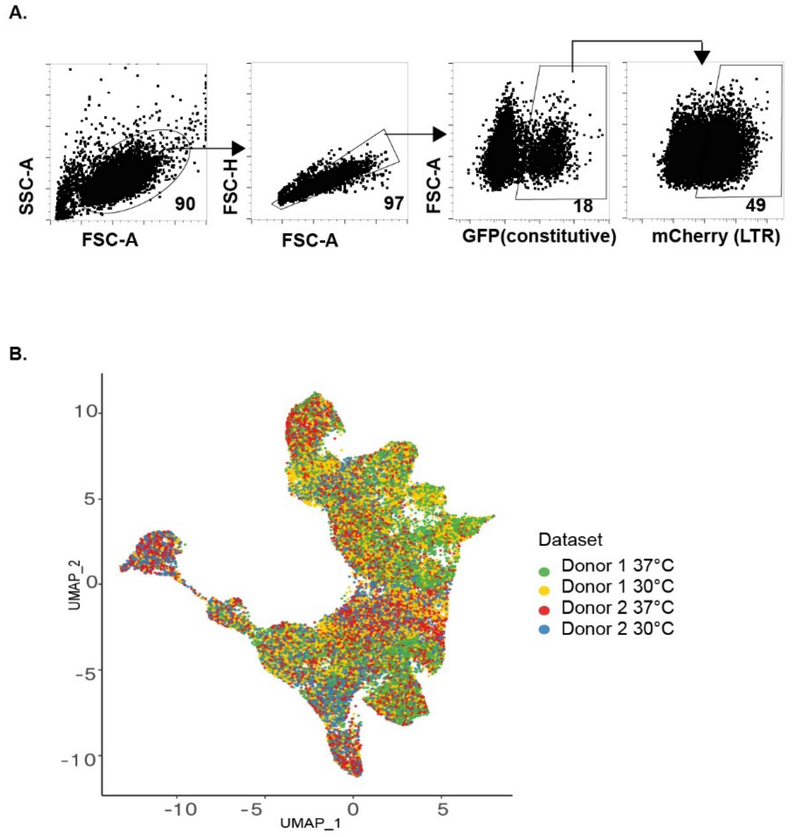

**Supplementary Fig. S1. Integration of single-cell data from different donors and conditions. A.** Gating strategy used to identify latently infected cells (GFP-positive, mCherry-negative) and actively infected cells (GFP-positive, mCherry-positive). The two right plots are the same as those shown in Fig. 1D. The arrow above the flow plots indicates the cell population gated and further analyzed in the subsequent plot. **B**. UMAP visualization of HSPCs cultured at the indicated temperature and transduced with HIV VT1. Cells were sorted for active or latent infection, pooled, and analyzed by single-cell RNA sequencing (see Fig. 1C for workflow). Data integration was performed using LIGER. Each point on the UMAP represents an individual cell, colored according to donor and culture condition.

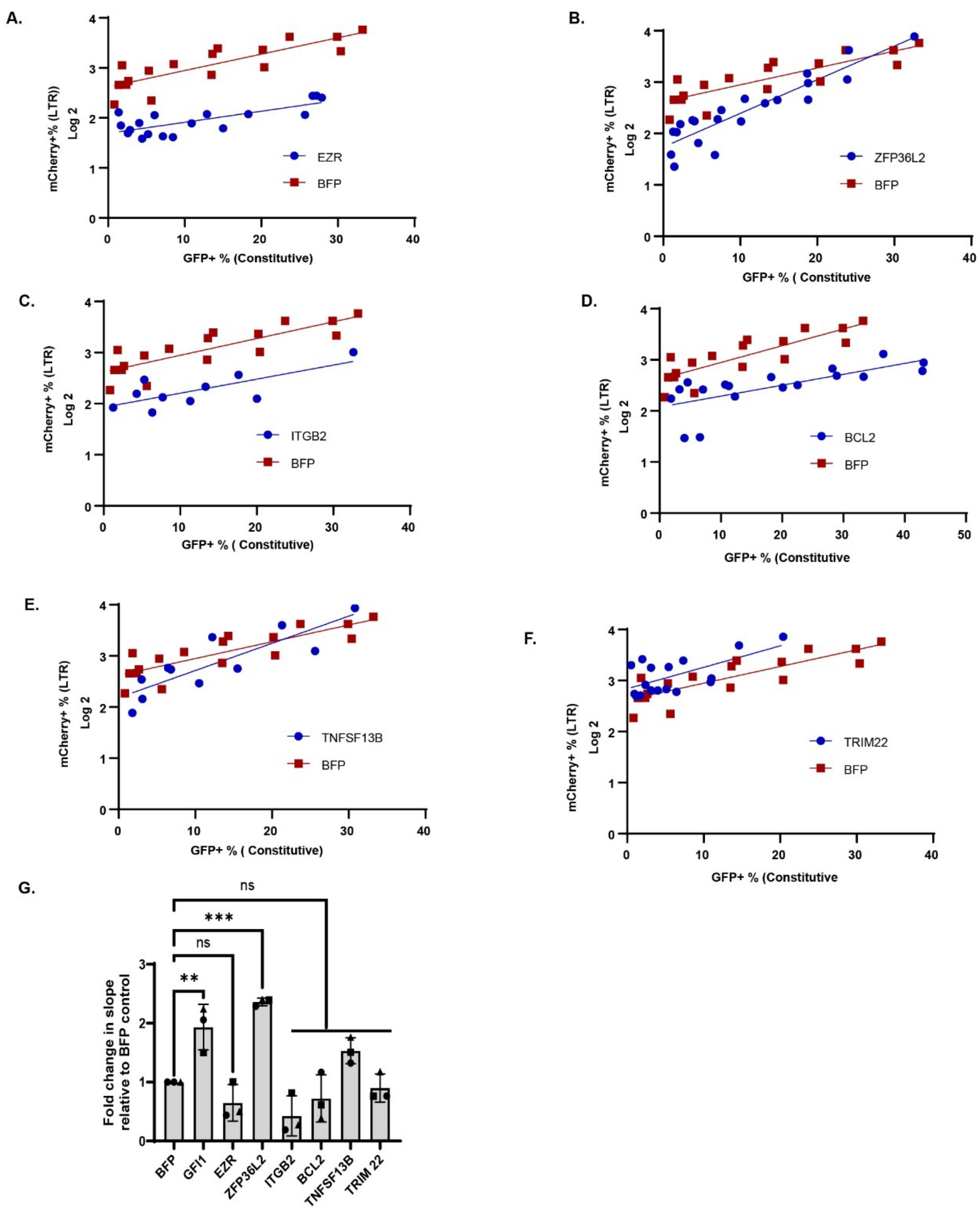

**Supplementary Fig. S2. Effect of the candidate latency-associated factors on HIV LTR-driven gene expression**. (**A-F**) Summary graphs showing the effect of the indicated factor on viral expression (mCherry) when cloned into VT1-IRES as in Fig. 3A and expressed in CEM-A2 cells as in Fig. 3C. **G.** Summary graph showing the fold change in slope obtained from using Deming regression (Model II) for the candidate genes. Fold change was determined by dividing the slope for each cDNA by the slope obtained for the control virus harboring BFP. Data are shown as mean ± standard deviation for *n*=3 independent experiments. Statistical analysis was performed using one-way ANOVA followed by Dunnett's multiple comparisons test. ***$P \leq 0.001$, **$P \leq 0.01$.

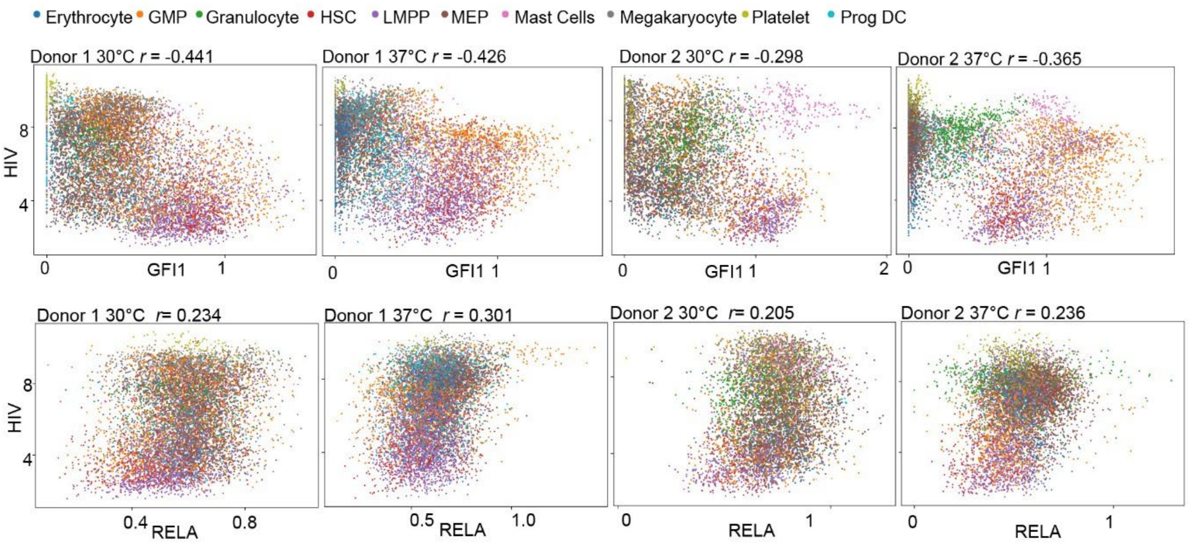

**Supplementary Fig. S3. GFI1 expression negatively correlates with HIV gene expression in primary HSPCs.** Correlation plots from the 4 scRNA-seq samples described in Fig. 1, illustrating the relationship between HIV gene expression and GFI1 or RELA expression levels.

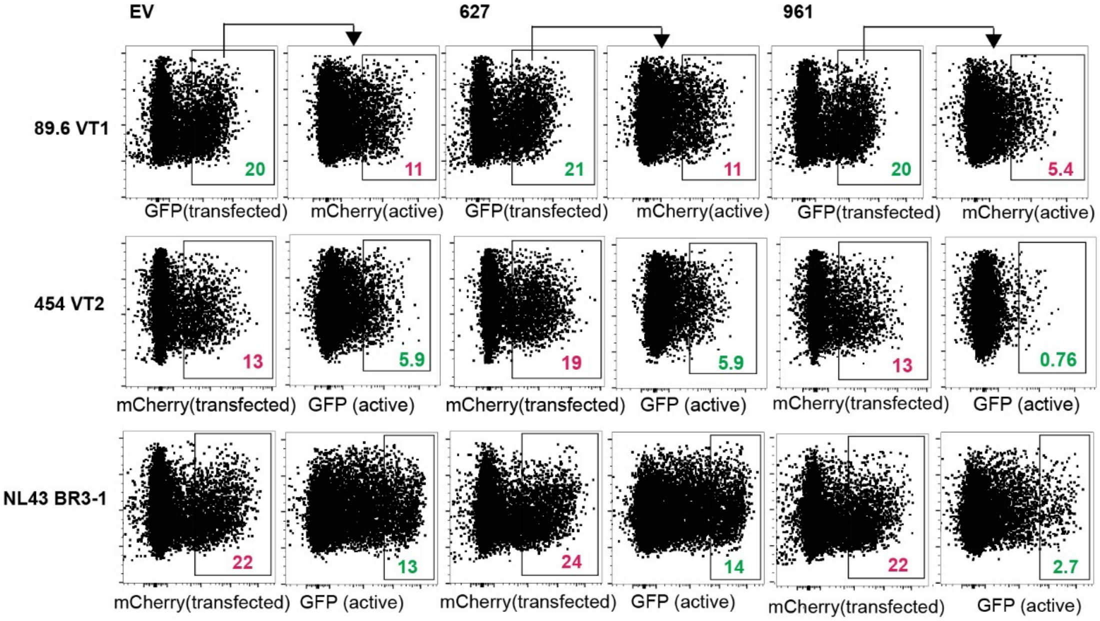

**Supplementary Fig. S4. Selective inhibition of HIV gene expression by Kruppel-associated box (KRAB) zinc finger proteins 961 versus 627 on HIV gene expression**. Flow cytometric analysis of HEK293T cells co-transfected as in Fig. 5A with the indicated HIV reporter constructs (VT1, VT2, or BR3-1; see Figs. 5B, E, H) and zinc finger protein. Corresponding data using GFI1 as the repressor are presented in Fig. 5C, F, I. The EV column is also included in Fig. 5 for direct comparison with the GFI1 condition.

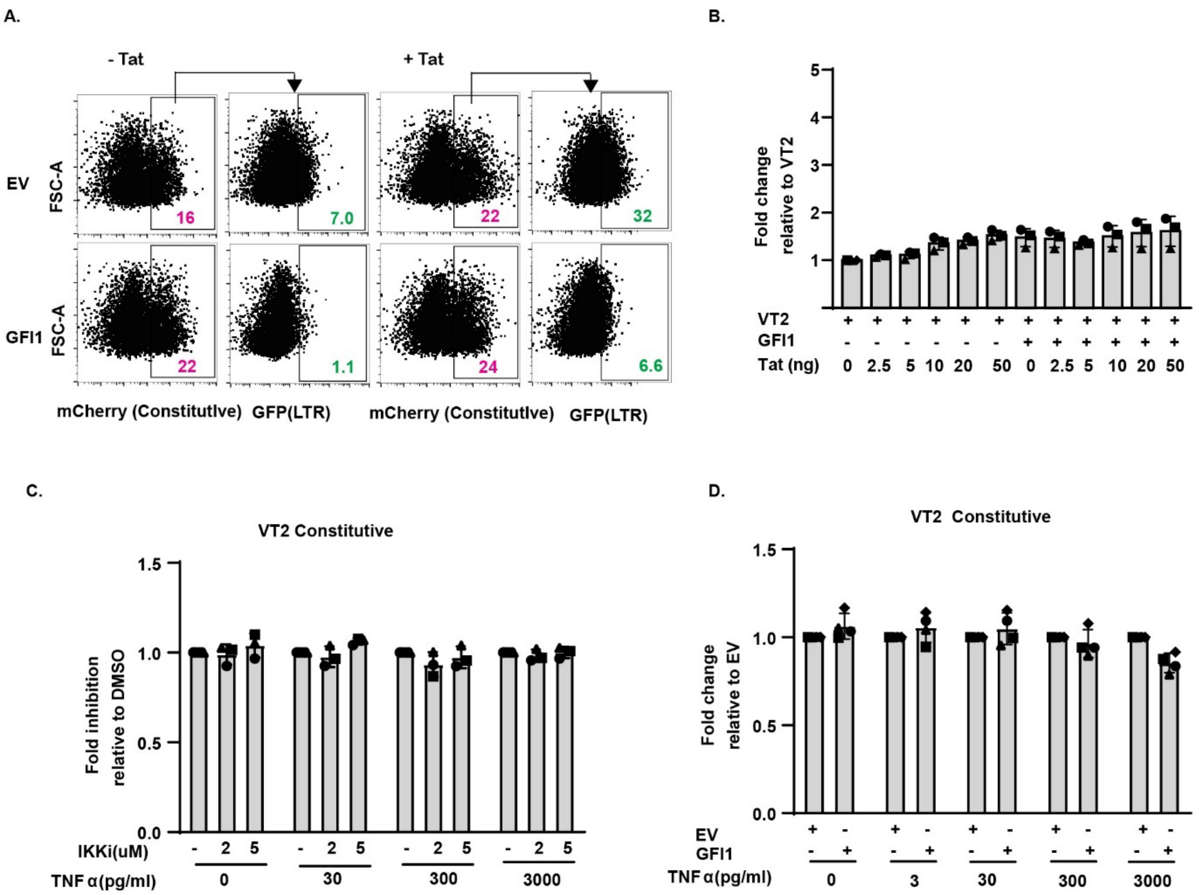

**Supplementary Fig. S5. Modulation of Tat and NF-κB activity alters LTR-driven HIV expression (GFP), but not constitutive promoter expression (mCherry) in VT2**. **A.** Representative flow cytometry plots of HEK293T cells co-transfected with the HIV reporter VT2, GFI1, and *tat*-expression plasmid as indicated in Fig. 7B. **B-D.** Summary graphs of constitutive promoter activity for experiments summarized in Fig. 7; corresponding LTR-driven activity shown in Figs. 7C, F, G, I respectively.

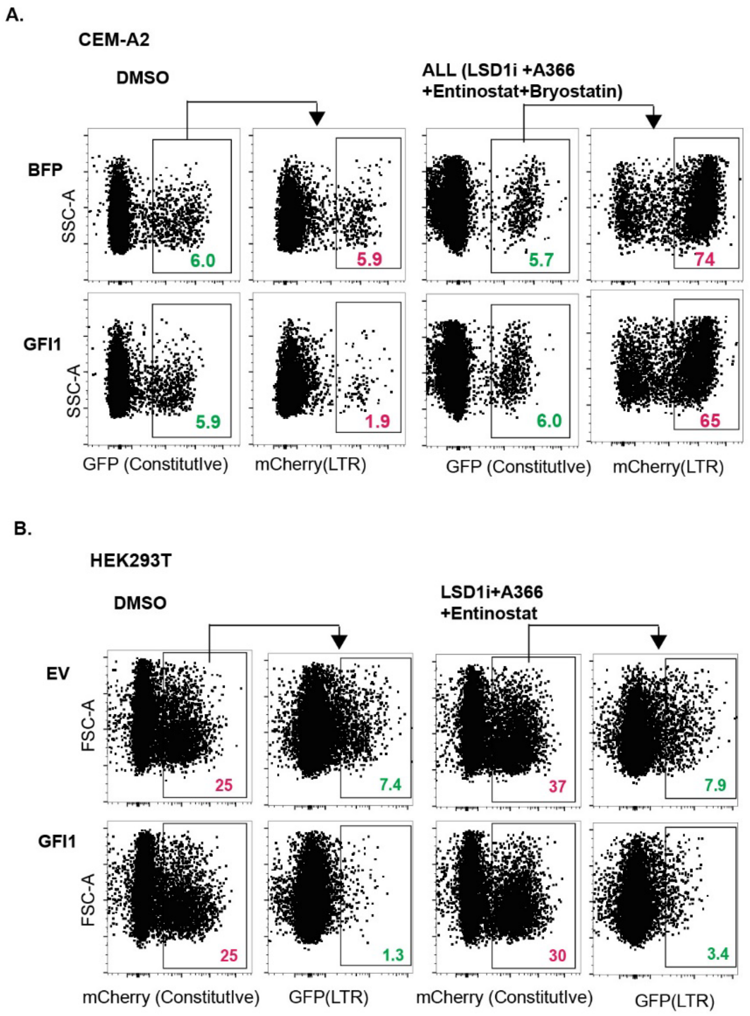

**Supplementary Fig. S6. A.** Representative flow cytometry plots of CEM-A2 cells transduced as in Fig. 8A, and treated with LRAs as indicated. **B.** Representative flow cytometry plots of HEK293T cells co-transfected as in Fig. 8D with VT2 and either pcDNA-EV or GFI1, followed by treatment with the indicated LRAs.

A.

| GFI1 | Y-intercept Fold Change(FC) |
|---|---|
| DMSO vs ALL | 35.1 |
| DMSO vs ENT+BRY | 30.8 |
| DMSO vs ENT | 13.8 |
| DMSO vs LSD1i+A366+ENT | 16.7 |
| DMSO vs BRY | 3.21 |
| DMSO vs LSD1i+A366+BRY | 4.39 |
| DMSO vs LSD1i | 1.15 |
| DMSO vs A366 | 1.22 |
| BFP | |
| DMSO vs ALL | 12.7 |
| DMSO vs ENT+BRY | 11.9 |
| DMSO vs ENT | 6.34 |
| DMSO vs LSD1i+A366+ENT | 7.61 |
| DMSO vs BRY | 2.19 |
| DMSO vs LSD1i+A366+BRY | 3.54 |
| DMSO vs LSD1i | 0.78 |
| DMSO vs A366 | 0.89 |

**Supplementary Table S1. Latency reversal agent (LRA)-dependent changes in slope and y-intercept, plus and minus GFI1 expression**. Table summarizing the effects of the indicated LRAs on HIV expression in the presence of GFI1 or BFP control. For each treatment, the fold change in the y-intercept was calculated from Deming linear regression (Model II) fitted to virus titration curves, where y-values were log2-transformed prior to analysis. ENT, entinostat; BRY, bryostatin; LSD1i, lysine demethylase inhibitor GSK-LSD1.
